# Boredom and the representation of information content in the neocortex

**DOI:** 10.64898/2026.07.09.737454

**Authors:** Johannes P.-H. Seiler, Jens-Bastian Eppler, Saman Seifpour, Lukas C. Wiese, Til Ole Bergmann, Florian Müller-Dahlhaus, Oliver Tüscher, Simon Rumpel

**Affiliations:** Institute of Physiology, Focus Program Translational Neurosciences (FTN), University Medical Center of the Johannes Gutenberg University Mainz, Duesbergweg 6, 55128 Mainz, Germany; Centre de Recerca Matemàtica, Edifici C, Campus Bellaterra, 08193 Bellaterra, Spain; Neuroimaging Center (NIC), Focus Program Translational Neurosciences (FTN), University Medical Center of the Johannes Gutenberg University Mainz, Mainz, Germany; Leibniz Institute for Resilience Research (LIR) GmbH, Mainz, Germany; Department of Psychiatry and Psychotherapy, University Medical Center of the Johannes Gutenberg University Mainz, Mainz, Germany; Department of Psychiatry, Psychotherapy and Psychosomatic Medicine, University Medicine Halle, Germany

## Abstract

Boredom – a pervasive mental state – promotes the pursuit of novel information by assigning negative value to monotonous conditions. Yet, how the brain extracts and represents the information content of ongoing sensory experience remains poorly understood. Here, we combine behavioral assays, neurophysiological recordings and computational modeling across humans and mice to investigate how sensory information shapes boredom-related behavior. In a cross-species choice task, both humans and mice robustly avoid monotonous sources of sensory stimulation. We formalize perceived monotony using empirical entropy as a measure of information content and show that monotony avoidance scales directly with low entropy and in humans correlates with boredom experience. Human electroencephalography and mesoscopic calcium imaging in mice reveal that the recruitment of neocortical activity tracks stimulus entropy. Two-photon calcium imaging in the auditory cortex of mice further uncovers a stimulus-invariant population code for entropy, supported by neurons tuned to information content. A recurrent network model reproduced this code through an interplay of afferent depression and recurrent facilitation. Together, we demonstrate how the information content of sensory experience is represented in cortical population activity, providing a basis for boredom-related avoidance behavior. Thus, our findings link synaptic and neuronal dynamics to boredom, acting as a safeguard mechanism to ensure high information input to the brain.

## Introduction

Boredom is a ubiquitous human experience, intimately familiar to many of us, arising across diverse everyday situations such as in school or workplace environments, particularly under conditions of monotony or insufficient stimulation^1–5^. Boredom is commonly defined as an aversive state of non-optimal mental engagement in which the demands of a situation fail to match an individual’s cognitive capacities^6–8^. Beyond its subjective unpleasantness, boredom has substantial clinical relevance, where elevated and chronic experiences of boredom are associated with a range of psychiatric conditions, including depression, substance use disorders, and attention-deficit disorders^9–15^.

Despite its broad psychosocial relevance, scientific research on boredom is comparatively young. Diverse efforts have been made to quantify boredom in humans, primarily utilizing self-report instruments such as the Multidimensional State Boredom Scale^16^ (MSBS), which captures momentary fluctuations in experienced boredom^17,18^. Experimentally, boredom is typically induced through prolonged exposure to sparse or repetitive sensory stimulation^19–22^, for example by presenting humans with intuitively boring videos of mundane activities such as hanging laundry^23^. Although such paradigms possess strong face validity, the specific environmental features that induce boredom remain poorly defined and lack a quantitative description.

While at a phenomenological level, boredom is characterized by negative valence, dissatisfying arousal, and a subjective slowing of time^6,24,25^, it is also thought to serve an important adaptive function^18,26–28^: By signaling odd situations such as monotonous environments, boredom generates a strong urge to disengage from the current situation and seek alternative sources of stimulation or activity^26,29,30^. As such boredom is believed to play an important role to safeguard constant information input to the brain, which is crucial for continuous learning, behavioral flexibility^31–37^, and prevents individuals from stagnancy in monotony^38,39^. This perspective aligns with broader “informavore” theories of cognition^31,40^, proposing that organisms are intrinsically motivated to seek information and maintain an optimal level of informational engagement^34,41,42^.

The idea that the nervous system is sensitive to information content is supported by several lines of evidence. Neuronal responses throughout sensory systems adapt and decline during repetitive stimulation, whereas rare or unexpected stimuli evoke enhanced neural responses^43–50^. Moreover, information-theoretic approaches are widely used in neuroscience to quantify how much information about external stimuli is encoded in neuronal activity patterns^51–53^. However, these approaches typically characterize information from the perspective of an external observer and do not explain how the brain itself might construct an internal representation of the information content of its sensory input as such. A translational framework that operationalizes boredom in terms of quantifiable information content across species, thus enabling experimental access to the underlying neural computations would therefore provide a critical step towards a better understanding of this intriguing mental phenomenon. While boredom-like behaviors have been described in several non-human species^54–60^, existing paradigms do not allow precise control or formal quantification of the information content available to the animal.

We here test the hypothesis that boredom reflects a mechanism to sense insufficient information and ask: How does the brain abstract, measure and represent the information content of ongoing sensory experience?

To this end, we leverage a recently developed behavioral choice paradigm reliably inducing boredom in humans through repeated sensory stimulation, while providing a simultaneous behavioral readout of boredom-driven avoidance behavior^22^. Taking advantage of the non-verbal nature of the task, we extend this paradigm and measure boredom in both humans and mice as behavioral avoidance of sensory sources with low information content. Using information theory, we quantify experienced information content as empirical Shannon entropy^61^ and investigate the representation of entropy in cortical activity, using EEG recordings in humans and mesoscopic calcium imaging in mice. In mice, we further examine the neural representation of information at the level of single neurons and neuronal populations using two-photon calcium imaging. We find that a representation of information content emerges from population dynamics that is largely orthogonal to stimulus identity. Finally, we employ a network model to identify computational mechanisms capable of explaining the observed neural dynamics. With this, we establish a translational and mechanistic framework for studying boredom across species and identify mechanisms that endow neuronal networks with the ability to abstract information content from sensory input.

## Results

### Avoidance of low sensory information content reflects experienced boredom in humans

To investigate how sensory information content shapes boredom in humans, we adapted a recently developed behavioral paradigm^22^. A cohort of 92 young adult, healthy subjects (see Supplementary Table 1 for demographic details) participated in a choice task, requiring them to choose between two buttons on a screen. Unbeknownst to the subjects, the two buttons represented either a *monotonous* alternative, leading to the presentation of a single repeated sound stimulus, or, a *variable* alternative, leading to the presentation of varying sound stimuli randomly drawn from a larger stimulus library (Figure 1a, Methods). Sound stimuli consisted of recordings of single spoken words of neutral meaning (Methods). As we were interested in the impact of the information content on choice behavior, rather than sound identity, we randomized the assignment of stimuli to the two alternatives across individuals and tasks. Each task ended after a defined number of trials and was free from reinforcement, apart from the auditory stimulation per se. In addition to the task comparing monotonous versus variable stimulation, participants also completed control tasks with both alternatives being monotonous or variable, thus providing equivalent stimulation. We used intermittent psychometric sentiment ratings on visual analog scales to assess the development of individually perceived boredom, affect and arousal during the task.

**Figure 1.**
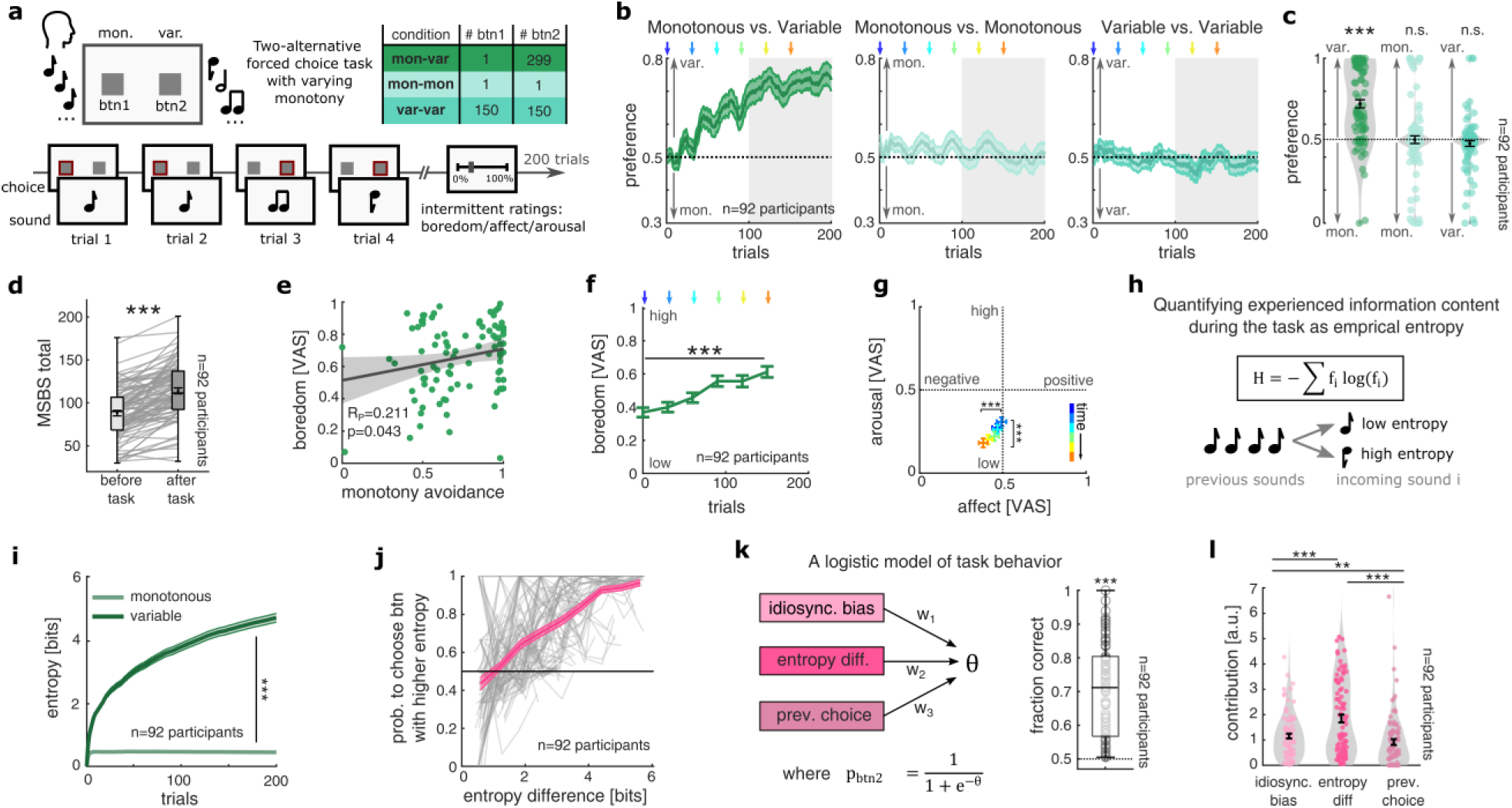
Individual monotony avoidance in humans reflects boredom and is driven by low information content: (**a**) Modified two-alternative forced choice task to assess human choice behavior between a monotonous alternative – a button (btn) providing repetitive stimuli – and a variable alternative – a button providing diverse stimuli (mon-var)^22^. Participants also completed control conditions with equivalent stimulation of both alternatives (mon-mon, var-var). Visual analog scale (VAS) ratings of current sentiment were collected intermittently. Table depicts stimulus library sizes for various conditions. (**b**) Mean preference over time across conditions (n=92 participants throughout). Arrowheads indicate ratings; data are mean ± SEM. (**c**) Distribution of mean preference during the second half of each condition (gray shading in b). Black bars, mean ± SEM; ***: p<0.001, n.s.: p>0.05. (**d**) State boredom before and after the choice task, assessed with the Multidimensional State Boredom Scale (MSBS). ***: p<0.001. (**e**) Correlation of mean perceived boredom and monotony avoidance (preference for the variable alternative) in the mon-var condition. Black line, linear fit; shading, 95% confidence interval. (**f**) Boredom ratings increased over the mon-var condition in parallel with monotony avoidance. Data are mean ± SEM; ***: p<0.001. (**g**) Affect and arousal ratings across the mon-var condition, indicating increasing displeasure and decreasing arousal over time. Data are mean ± SEM; ***: p<0.001. (**h**) Schematic of empirical entropy as a trial-wise measure of experienced information content (Methods). (**i**) Mean experienced entropy for the monotonous and variable alternatives over time in the mon-var condition. Data are mean ± SEM; ***: p<0.001. (**j**) Choice probability in the task as a function of difference in previously experienced entropy between both alternatives. To compute the choice probabilities, the single-trial data was binned according to experienced entropy difference (Methods). Gray lines, individual participants; pink line and shading, mean ± SEM. (**k**) Logistic regression model incorporating sensitivity to experienced entropy differences, idiosyncratic choice bias and a tendency to repeat the previous choice (left), and individual prediction performance (right; median >70%; ***: p<0.001). (**l**) Contributions of model parameters to choice behavior, showing a dominant contribution of entropy; **: p<0.01, ***: p<0.001.

Over the course of approximately 100 trials, participants developed a bias avoiding the monotonous alternative over the variable alternative (Figure 1b,c; for full statistical details see separate Statistical Information file). In task conditions with equivalent stimulation, subjects did not show a bias to either side. The procedures led to an overall increase of boredom scores on the Multidimensional State Boredom Scale (MSBS), a standardized psychometric questionnaire widely used in boredom research (Figure 1d, Supplementary Figure 1a). Corroborating previous observations^22^, the extent of monotony avoidance in individual participants was positively correlated with the mean boredom ratings during the task, indicating a reflection of perceived boredom in choice behavior (Figure 1e). Boredom ratings gradually built up over the course of the task, rising in parallel to the increasing choice bias of avoiding the monotonous alternative (Figure 1f). This was followed by gradually decreasing ratings of affect and arousal (Figure 1g), adhering to classical definitions of boredom as an unpleasant affective state of low arousal^6,25,62,63^. In the control conditions, the dynamics of sentiment across time were less pronounced and showed highest levels of boredom for the condition of full monotony (Supplementary Figure 1b-d).

We next quantified the empirical Shannon entropy^61^ within the sequence of sound stimuli sampled from each alternative until a given trial, relative to all sampled stimuli so far, providing a measure of experienced information content (Figure 1h, see Supplementary Figure 1e and Methods for details). As expected, entropy sampled from the variable alternative gradually built up over the trials in the task, whereas it remained at a low level for the monotonous alternative (Figure 1i). To assess how the differences in experienced entropy between both alternatives up to a given trial would affect the future choice of participants, we binned the data according to the entropy difference in each trial, and analyzed the upcoming choice probability for the alternative providing higher entropy (Figure 1j). Interestingly, we found that higher entropy sampled from one alternative predicted a higher probability of sampling this alternative again, suggesting that preference behavior in the task was driven by avoidance of low information content. To investigate this aspect in greater detail, we used logistic regression to model individual choice behavior on a single-trial basis and compared the contribution of sampled entropy to other choice factors. Specifically, we used three regressors: (i) The sampled *entropy difference*, providing a measure of how sensitively a subject adjusts its choice to experienced information content, (ii) an *idiosyncratic choice bias*, describing an inherent, stimulus-independent preference towards a given side^64,65^, and (iii) the *previous choice*, describing behavioral inertia to repeat the previous choice independent of the sampled stimuli^66–68^ (Figure 1k, Methods). The model captured a significant fraction of variance in decision behavior, predicting approximately 70% of all choices correctly. Interestingly, comparing the contribution of all regressors (Methods), we found that entropy difference showed the highest impact on choice behavior (Figure 1l).

Together, these observations indicate that boredom in humans is directly driven by low sensory information content, triggering avoidance of sensory sources associated with low information content.

### Mice avoid sensory sources with low information content alike bored humans

In order to gain insight into the mechanistic underpinnings of boredom in an experimentally accessible model organism, we devised a place preference task that analogously prompted mice to choose between monotonous or variable auditory stimulation under conditions of low arousal. Again, individual choice behavior was uncoupled from any feedback except the sensory stimulation per se. In this task, conducted with a sample of 24 wildtype C57BL/6J mice (see Supplementary Table 2), the position of a given mouse was continuously tracked inside a Skinner box which was divided into two zones, either coupled to the presentation of monotonous or variable sound stimuli at an interval of 3 seconds (Figure 2a, Methods). The stimulus set consisted of diverse naturalistic and artificial sounds of various lengths (70-1,000ms) designed not to carry overt ethological relevance (see Methods for details). Analogous to the human tasks, we randomized the assignment of stimuli to the two zones. In addition to the comparison between monotonous and variable stimulation, mice underwent various control conditions, comprising equal types of stimulation in both zones analogous to the human experiment. In addition, we conducted control experiments involving no auditory stimulation at all (*silence*) or the specific emission of loud white noise (*noise*) previously reported to elicit avoidance responses^69,70^.

**Figure 2.**
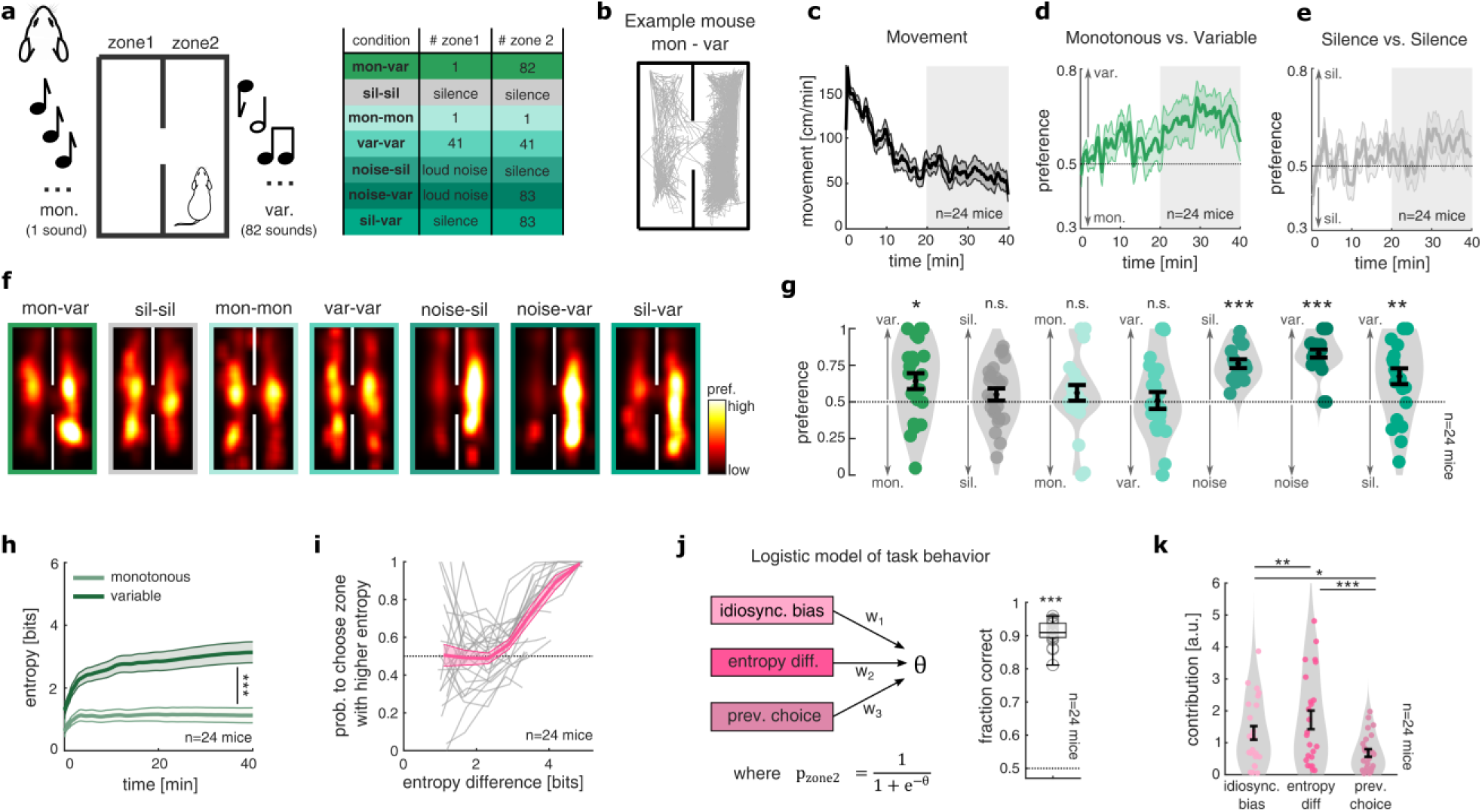
Mice exhibit analogous monotony avoidance driven by low information content: (**a**) Translational mouse version of the choice task between monotonous and variable auditory stimulation. Mice were tracked in Skinner box with two mirror-image zones and presented with different types of auditory stimuli depending on their current location (left). In addition to the comparison of monotony and variability (mon-var) and equivalent stimulation (mon-mon, var-var), mice underwent various control conditions including no stimulation (sil: silence) or loud white noise (noise) (right). Table depicts library sizes and stimulus types for various conditions. (**b**) Movement trajectory of one example mouse in a session of the mon-var task. (**c**) Mean movement over time in the mon-var condition (n=24 mice throughout), showing progressive settling within one zone. Data are mean ± SEM. (**d,e**) Mean place preference over time in the mon-var (d) and silence (e) conditions. Mice avoided monotonous stimulation but showed no bias between silent zones. Data are mean ± SEM. (**f**) Mean spatial occupancy across mice and sessions for all conditions. (**g**) Mean place preference during the second half of each condition (shading in d,e), showing preference for less monotonous stimulation and avoidance of loud white noise. Black bars, mean ± SEM. (**h**) Mean experienced empirical entropy for the monotonous and variable zones over time in the mon-var condition (Methods; Figure 1h,i). Data are mean ± SEM.; ***: p<0.001. (**i**) Choice probability as a function of the difference in previously experienced entropy between zones. Data were binned by entropy difference (Methods). Gray lines, individual mice; pink line and shading, mean ± SEM. (**j**) Logistic regression model incorporating sensitivity to experienced entropy differences, idiosyncratic choice bias and choice repetition (left), and prediction performance for mouse location (right; >90% correct; ***: p<0.001). (**k**) Contributions of model parameters to choice behavior, showing a dominant contribution of entropy, consistent with the human results (see Figure 1l). *: p<0.05, **: p<0.01, ***: p<0.001.

In a given task session, mice typically explored both zones of the box (Figure 2b), showing high amounts of movement before settling down in one of the zones (Figure 2c). Interestingly, in the sessions choosing between monotonous and variable stimulation, mice also exhibited a systematic bias avoiding the monotonous alternative (Figure 2d). This bias built up over time, reaching a plateau after approximately 20 minutes, resembling monotony avoidance in human participants. Moreover, we found that differences in monotony avoidance of individual mice remained comparably stable across sessions, suggesting a link with individual trait properties (Supplementary Figure 2a,b). In control conditions with equivalent stimulation in both zones, no systematic bias emerged, however, mice showed avoidance of loud white noise as expected (Figure 2e, pooled preferences are shown in Figure 2f,g). Interestingly, mice even showed a preference for variable sound stimuli as compared to silence, suggesting an overall seeking for sensory stimulation^56^. Avoidance biases in asymmetric control conditions showed a similar development within a session as in the task comparing monotonous versus variable stimulation, and were largely stable across sessions (Supplementary Figure 2c,d). Moreover, the overall levels of movement and switching between the zones were comparable across task conditions (Supplementary Figure 2e,f).

In analogy to our human experiments, we quantified information content by estimating the empirical entropy associated with the two alternatives of sensory stimulation. We split the continuous session data into a trial structure according to the stimulus presentation intervals (Methods) and quantified the empirical entropy sampled from either zone to obtain a measure of experienced information content. In line with our human data, we found that mice on average accumulated increasing levels of entropy by sampling the variable zone, whereas the monotonous zone was associated with low levels of entropy throughout (Figure 2h). Interestingly, when analyzing choice probabilities for individual mice as a function of the difference in sampled entropy from the two alternatives, we observed that higher entropy for a given alternative was associated with higher preference, similar to our findings in humans (Figure 2i, compare to Figure 1j), albeit the probability curves of mice, on average, started to rise at higher levels of entropy (human: ∼1.5 bits, mouse: ∼3 bits). Applying an equivalent logistic regression model as in our human analysis to quantify the effects of *entropy difference*, *idiosyncratic choice bias* and *previous choice* on mouse behavior, we found that the model predicted a high fraction of place preference correctly (∼90%, Figure 2j), revealing a similar contribution pattern for the three regressors with experienced entropy having the highest association with decision behavior (Figure 2k).

These observations indicate that mice reliably avoid sensory sources of low information content, resembling boredom-related behavior in humans.

### Information content is reflected by increased cortical recruitment and arousal

Our assays in humans and mice revealed an essential role of experienced entropy in explaining boredom-related avoidance behavior, suggesting that mice and humans share an internal representation of information content to guide affective and behavioral responses. We next sought to investigate in how far information content is reflected in neurophysiological signatures observable in both species.

To this end, we combined assessments of overall cortical activity while presenting individuals with sensory stimuli that systematically varied in information content. In addition, we used pupil size measurements to obtain a physiological readout of arousal^71,72^, motivated by the observed linkage between experienced boredom and low degrees of arousal in humans^25^. Various previous studies have demonstrated a robust modulation of sound-evoked responses in the auditory cortex by the statistics and predictability of sound events, classically described under terms such as “adaptation”, “repetition suppression”, or “mismatch negativity”^44,45,47–49,73–75^. While these studies often use categorical comparisons of responses for expected standard stimuli versus unexpected deviant stimuli, the neural signatures of different degrees of expectedness on a continuum remains less understood. To address this aspect, we used a library of ten neutral, but distinctly represented sound stimuli as a basis to create a set of sequence stimuli systematically varying in information content (see Methods). We constructed 20 different sequence patterns with varying degrees of repetitiveness, each comprising 10 sound stimuli presented at inter-sound intervals of 1 second. This design allowed us to quantitatively describe the information content conveyed by the statistical structure of the sequence up to a given stimulus by computing entropy (Figure 3a). While various types of entropy have been proposed, capturing different aspects of stimulus novelty and surprise^76^, we quantified the basic Shannon entropy congruent to the information measure that explained well the behavioral choice bias (Supplementary Figure 3, Methods). In order to control for response components that are driven by the physical differences of the stimuli, we constructed different versions of a particular sequence pattern by varying the composition of sound stimuli. Sound sequences for human and mouse experiments had the same structure, but were constructed from different sound libraries matching the respective hearing range, and contained 3 versions (human, i.e. 60 sequences total) or 5 versions (mouse, i.e. 100 versions total) of each sequence pattern (see Supplementary Figure 4).

**Figure 3.**
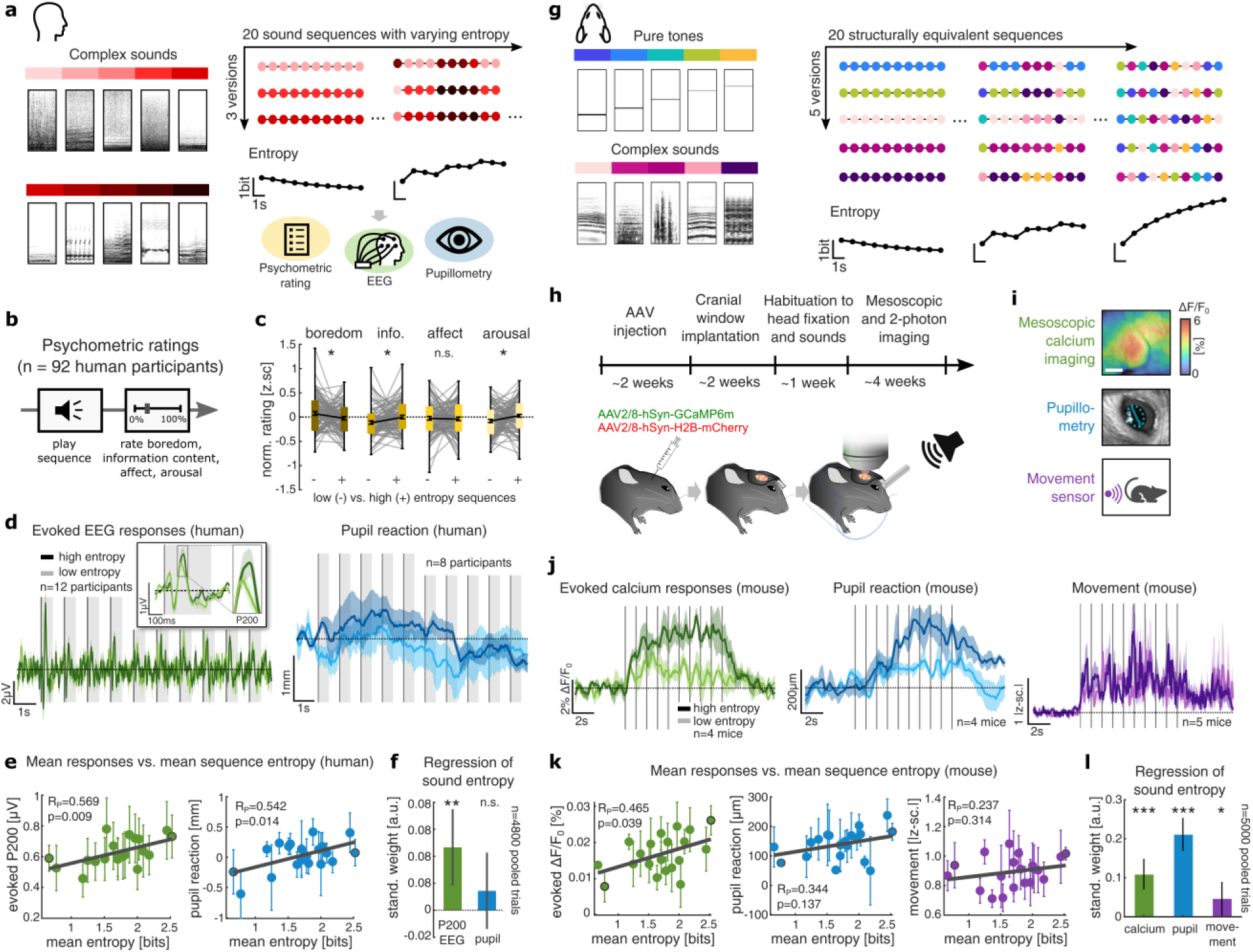
Sensory information content is reflected by the magnitude of evoked cortical activity in humans and mice: (**a**) Ten sounds (spectrogram range, 16 Hz–15 kHz, 0.5s) were used to generate 20 entropy-varying sequences of ten sounds, each presented in three stimulus versions (60 sequences total). Participants either rated the sequences (n=92) or underwent 64-channel EEG and pupillometry during passive listening (n=12). (**b**) After each sequence, participants rated perceived boredom, information content, pleasantness (affect) and arousal. (**c**) Ratings for the five highest- and five lowest-entropy sequences (n=92). Box plots show median and interquartile range; vertical bars, mean ± SEM; *: p<0.05, n.s.: p>0.05. (**d**) Mean EEG responses across Fz and Cz (left) and pupil responses (right) to example high- and low-entropy sequences. Inset, mean sound-evoked potentials across all sounds in each sequence, showing a larger P200 for the high-entropy sequence (see Supplementary Figure 5b,c; EEG responses are from n=12 participants; pupil responses are from n=8 participants). Data are mean ± SEM; dark and light traces indicate high- and low-entropy sequences, respectively; gray shading indicates stimulus presentation and dashed lines baseline. (**e**) Correlations of mean entropy and mean evoked P200 amplitude (left), or pupil dilatation (right) across 20 sequences, averaged across stimulus versions and participants. Gray lines show linear fits; error bars, SEM. (**f**) Standardized coefficients from a linear regression relating P200 and pupil responses to mean sequence entropy. Error bars, 95% confidence intervals. n.s.: p>0.05, **: p<0.01. (**g**) Analogous mouse experiment using the same 20 sequence structures in five stimulus versions (100 sequences total), generated from five pure tones and five complex sounds of 70ms duration (n=5 mice). (**h**) Experimental preparation for mesoscopic and two-photon calcium imaging. (**i**) Mesoscopic imaging of the auditory cortex was combined with pupillometry and movement assessment during passive sequence presentation. (**j**) Mean calcium (left), pupil (center) and movement (right) responses to example high- and low-entropy sequences. Data are mean ± SEM; dark and light traces indicate high- and low-entropy sequences, respectively. Dashed lines represent baseline mean. (**k**) Correlations between mean sequence entropy and evoked calcium activity, pupil responses and movement, as in e. (**l**) Standardized regression coefficient, as in f, showing associations of sequence entropy with evoked activity and pupil responses. Error bars, 95% confidence intervals; *: p<0.05, ***: p<0.001.

To validate our stimulus set with respect to evoked sentiment, we collected psychometric ratings from 92 human participants (same sample as for the behavioral experiment in Figure 1, see Methods), measuring the perceived levels of *boredom*, *information content*, *affect* and *arousal* for each sequence (Figure 3b). When comparing mean sentiment for the quartile of sequences with highest average entropy, against the quartile with lowest average entropy, we found that the high-entropy sequences evoked significantly less boredom, higher degrees of perceived information and higher arousal than the low-entropy sequences, despite the overall highly arbitrary character of the sound sequences (Figure 3c). Correlating the sentiment across the full set of sequences with mean entropy, showed analogous effect trends (Supplementary Figure 5a). For rated levels of affect, we did not observe a significant modulation with this assay. Together, these observations corroborate that varying complexity in sequence structure, quantified by entropy, induces systematic differences in perception and sentiment.

In a next step, we recruited a new cohort of twelve human participants and recorded electroencephalographic (EEG) activity of the auditory cortex alongside pupil responses during presentation of the sound sequences (see Supplementary Table 3, Methods). Both evoked EEG and pupil responses are physiological markers previously linked to surprise and novelty detection^43,44,48–50,77–83^. We sought to test in how far experienced entropy may be utilized as an essential, parametric feature of the stimuli allowing to predict these physiological responses. As it is well described that novel stimuli modulate early auditory event-related potential (ERP) components^44,84,85^, we analyzed the mean auditory evoked potentials in frontocentral midline electrodes and tested how strongly they were modulated by differences in entropy (Supplementary Figure 5b,c). We observed a pronounced modulation of responses with entropy in the P200 latency range, an ERP component previously linked to sensory expectations and salience detection^50,86,87^, so that we decided to further analyze this component as a correlate of varying information content. While the mean EEG response size across a session remained stable (Supplementary Figure 5d), sequences with high information content indeed induced more pronounced sound-evoked potentials and greater pupil dilation (Figure 3d). When averaging responses across all sounds and versions of the same sequence patterns, mean entropy correlated significantly with both, evoked P200 EEG potentials and pupil dilation (Figure 3e). At higher temporal resolution, linear regression of entropy associated with a given sound presentation within a sequence with the corresponding response amplitudes revealed a significant relationship for auditory P200 signals (Figure 3f, Supplementary Table 4). Also, pupil size showed a positive regression weight, but did not reach statistical significance, likely due to slower time dynamics (Figure 3f, Supplementary Table 4). These results indicate that the information content of sensory sequences is reflected in the recruitment of human auditory cortical activity and more pronounced pupil responses.

We then prepared five wildtype C57BL/6J mice for calcium imaging of sound-evoked responses in the auditory cortex, injecting them with adeno-associated virus (AAV)-based vectors to express GCaMP6m for activity imaging, and H2B::mCherry for structural marking of nuclei (Figure 3h, Methods). Before data acquisition, mice were extensively habituated to head fixation and pre-exposed to the sound stimuli. While presenting the sound sequences to the mice, we measured mesoscopic calcium activity^88^ and conducted simultaneous recordings of pupil size and movement (Figure 3i; see Supplementary Table 5 for details on the dataset). Resembling our observations in humans, we also observed that sequences with higher average entropy evoked more pronounced mesoscopic calcium responses, pupil responses, and stronger movement (Figure 3j). The mean mesoscopic response remained stable throughout a session (Supplementary Figure 5e). Across all sequence patterns and their versions, mean entropy correlated significantly with mean evoked mesoscopic calcium responses (Figure 3k). Pupil size and movement also showed a positive, but non-significant correlation with entropy. Regressing entropy from the response data to individual sounds revealed a positive association with all three measures, mesoscopic calcium response, pupil size, and – less pronounced – movement (Figure 3l, Supplementary Table 4).

Together, these observations in humans and mice show that information content of sensory input is reflected by the average evoked response in the auditory cortex and is associated with heightened physiological arousal.

### The representation of information content at the level of single neurons

The experimental translation of boredom-related neural processing to mice, facilitated accessing the representation of information content with single neuron resolution. In the five mice prepared for calcium imaging, we performed high-resolution two-photon calcium imaging after functionally identifying the tonotopic structure of the auditory cortex using intrinsic imaging (Supplementary Figure 6a,b; Methods). The awake mice were head-fixed and we recorded neurons in layer 2/3 of the auditory cortex while presenting the sound sequences in a pseudorandom order at a sound pressure level of 65 dB (Figure 4a). This dataset comprised 7,392 neurons in 44 fields of view (FOV) of which approximately 74%, i.e. 5,454 neurons, showed a significant response to at least one of the ten sounds (see Supplementary Table 6 for details on the dataset). The calcium responses of single neurons to the different sounds in the sequences revealed a diversity of response patterns (Figure 4b,c). For a given sound, on average ∼17% of neurons showed an excitatory response with increased ΔF/F_0_ signal, whereas ∼4% of neurons showed a suppressive response with decreased ΔF/F_0_ signal, aligning with the findings of previous studies^89–91^ (see Methods, see Supplementary Figure 6c-g for response types across stimuli).

**Figure 4.**
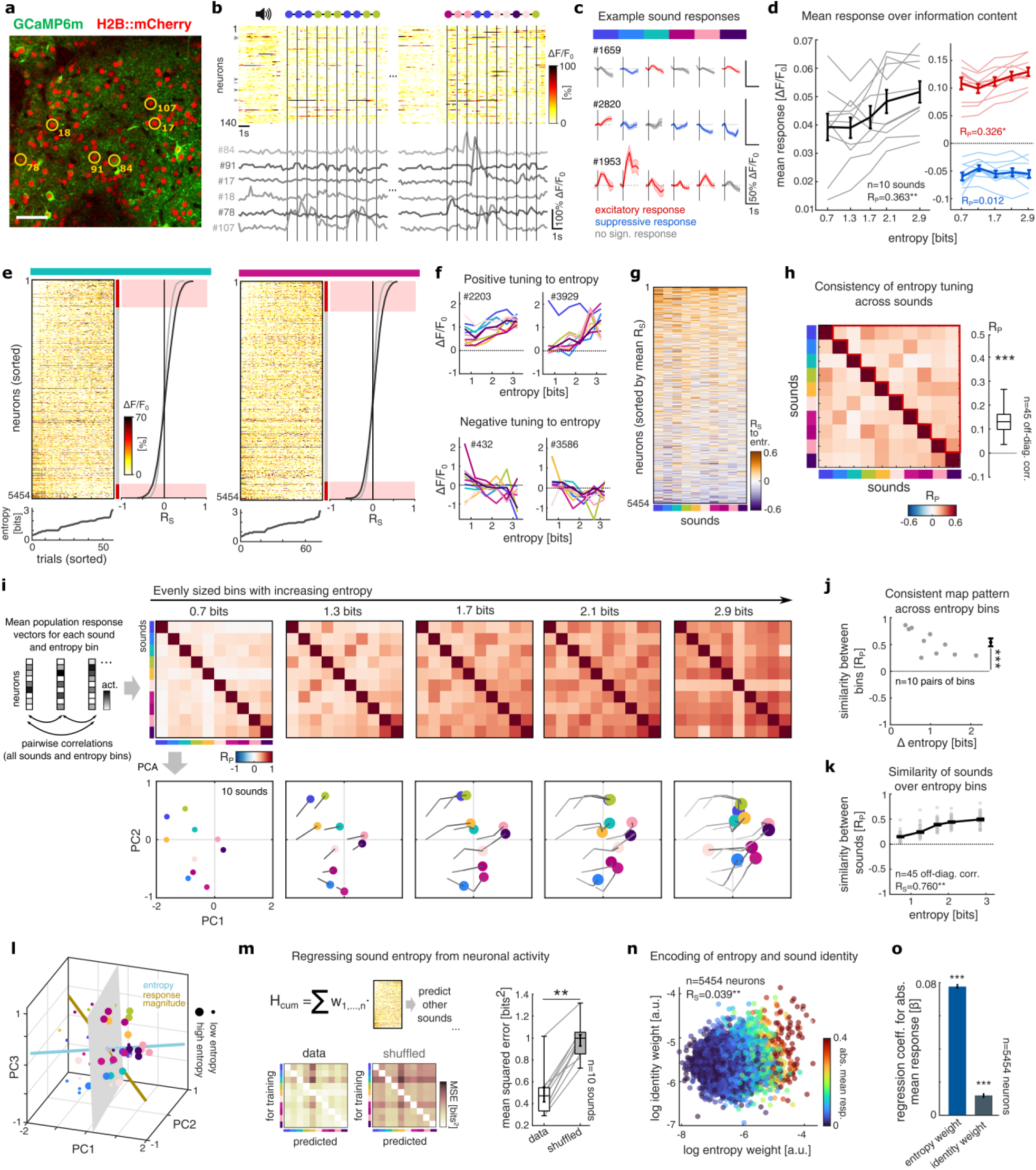
A representational code of information content driven by neurons tuned to stimulus entropy: (**a**) Example FOV from two-photon calcium imaging, showing broad GCaMP6m and H2B::mCherry expression in cortical layer 2/3 neurons. Scale bar: 50μm. (**b**) Calcium responses (ΔF/F0) to sound sequences with varying entropy of example neurons. (**c**) Mean ΔF/F0 responses to six example sounds (rows) for three example neurons (displayed as mean ± SEM). Responses are categorized into excitatory, suppressive or non-significant activities. (**d**) Mean overall sound-evoked activity over five evenly sized bins of entropy across sounds (left). The right panel shows the same analysis for excitatory (red) and suppressive (blue) responses. Gray lines represent different sounds. Thick lines, mean ± SEM. (**e**) Evoked activity of all sound-responsive neurons (rows) across presentations of an example pure tone (left, green) and complex sound (right, magenta). Trials (columns) are ordered by increasing entropy (below); neurons are ordered by decreasing Spearman correlation between activity and entropy. Correlations, reflecting entropy-tuned activity, are shown at right for observed and shuffled data; red bars mark significant correlations after Bonferroni-Holm correction. (**f**) Mean evoked activity of example neurons over entropy bins, showing positive tuning (top panels) and negative tuning (bottom panels) to entropy. Lines indicate different sounds. (**g**) Consistency of entropy tuning across sounds. Neurons (rows) are ordered by their mean correlation of activity and entropy across sounds; consistent gradients across columns indicate stimulus-invariant entropy tuning. (**h**) Pearson correlations between entropy tuning coefficients across sounds (left) and comparison with zero (right) (***: p<0.001). Box plots show median and interquartile range. (**i**) Representational similarity analysis of mean population responses to all sounds across five equally sized entropy bins. Pairwise correlations between population response vectors yielded representational similarity matrices (top). PCA projections (bottom) show representations of individual sounds; colors indicate sound identity, point size indicates entropy bin and gray lines connect representations of the same sound across lower-entropy bins. Increasing entropy systematically shifted sound representations along a common axis. (**j**) Correlations between off-diagonal representational similarity patterns for all pairs of entropy bins (black bar: mean ± SEM; ***: p<0.001). (**k**) Mean off diagonal representational similarities over entropy bins. The positive correlation reflects the increase in average representational similarity (see increasing of red color top panel of i). Black line displays mean ± SEM. (**l**) Three-dimensional PCA projection of sound representations across entropy bins. Point size indicates population responses to low- or high-entropy stimuli; the gray plane shows the linear decision boundary for entropy decoding. Lines indicate the best-fitting representational axes for entropy (light blue) and mean response magnitude (ochre), which were not fully aligned. (**m**) Cross-stimulus linear regression of entropy from single-trial population responses. Models trained on each sound were tested on all other sounds (bottom left); shuffled data are shown for comparison (bottom right). Population activity supported stimulus-invariant regression of entropy (right, **: p<0.01). (**n**) Correlation of the average entropy regression weights and identity decoding weights across neurons. Color indicates absolute mean response magnitude. (**o**) Regression coefficients for entropy weights and identity weights predicting absolute mean response magnitude. (error bars reflect 95% confidence intervals, ***: p<0.001).

For our further analyses we only considered the 5,454 sound-responsive neurons, pooled across all mice and FOVs. In our stimulus design, the presentation of a given stimulus was associated with a varying degree of information content, determined by the respective sequence context. We therefore sorted the single trial responses for each stimulus into five evenly sized bins of increasing entropy (see Methods). In line with our previous findings on the mesoscopic level, we observed that the mean response amplitude increased with growing entropy, largely driven by a rise in excitatory responses, whereas suppressive responses remained widely stable (Figure 4d).

To analyze the tuning of neurons to information content, we next constructed single-trial population response vectors based on the evoked calcium activity within a 600ms time window after stimulus presentation of all 5,454 neurons. Sorting the response vectors for all trials of a given sound by entropy, revealed that some neurons showed significant positive (∼13%) or negative (∼7%) tuning to entropy (Figure 4e). For individual neurons, positive or negative tuning to information content was similar when analyzing responses to different sounds (Figure 4f). To determine whether individual neurons exhibited shared entropy tuning across sounds, we computed the correlation of activity and entropy, for each neuron and sound (Figure 4g), and then calculated the pairwise correlations of the resulting tuning vectors across sound stimuli (Figure 4h). We observed a significant degree of similarity in the tuning of neurons across sounds, indicating that a fraction of neurons modulates its activity according to the information content independent of the identity of the incoming stimulus.

### The representation of information content at the level of neuronal populations

Going beyond the tuning of individual neurons to entropy, we next investigated how information content is encoded in the structure of population activity in the auditory cortex. To this end, we applied a representational similarity analysis (RSA), which provides an unsupervised estimation of the relational structure of the population responses evoked by the sound stimuli varying in entropy^92–95^. The trial-to-trial reliability of sound-evoked response patterns was highest in the bin with low entropy, and remained at lower, comparable levels in bins with higher entropy (Supplementary Figure 7a). We computed the average population response vectors for each sound and entropy bin, respectively, and generated a representational similarity matrix by computing the pairwise Pearson correlations (Methods). We then applied a principal component analysis (PCA) to the full representational similarity matrix in order to visualize the representational structure of sounds in dimension-reduced space^90,91,95^. For each of the five entropy bins, we displayed the representational similarity matrices and the respective projections in the shared PCA coordinate system (Figure 4i). Here, we made three main observations: First, the relative distances between the activity patterns evoked by the ten different sounds remained largely preserved in each of the five entropy bins. This was reflected by significant positive correlations when comparing the off-diagonal entries of the five representational similarity matrices (Figure 4j). Second, the mean pairwise similarities in a given similarity matrix increased with entropy (Figure 4k), indicating that sound-evoked response patterns with high information content were generally more similar. Third, sound representations displayed a systematic shift with growing entropy along a common axis, closely aligned to the first principal component (see trajectories in Figure 4i).

To investigate in how far the direction of shift with entropy in representational space was shared or independent for each sound, we estimated the axes that best captured entropy for each stimulus individually^91^ and compared their orientations (Methods). We found a significant alignment of representational entropy axes across sounds, indicating a stimulus-invariant representation of information content in cortical activity (Supplementary Figure 7b,c). Given various reports of strong adaptation of responses in the auditory cortex^44,45,96^ and our previous finding that entropy is reflected in the level of overall cortical activation, we wondered in how far the representational axis capturing entropy essentially reflects stimulus adaptation. We therefore estimated the overall representational axis for entropy in our dataset by considering all sound responses, as well as the representational axis that best explained differences in response magnitude (Figure 4l, Methods). We found that despite an overall positive correlation between both axes, they were statistically distinct and entropy-encoding was not explained by mere differences in mean responsiveness (observed data: R_P_=0.464, 95%-CI = [0.009 0.560]; Supplementary Figure 7d,e). This indicates entropy is not only encoded by a modulation of overall activity but additionally in the structure of evoked activity patterns.

The presence of a shared entropy axis for the various sounds tested, implies that the level of entropy conveyed by a particular stimulus can be decoded in a linear manner from the population activity, independent of the particular sound identity. To test this, we trained a binary decoder classifying responses into ‘high entropy’ or ‘low entropy’ bins and observed a performance level of approximately 86% correct classifications (Figure 4l). To further corroborate this, we trained a linear regression model to predict entropy from the single-trial population response vectors of one given sound, and applied this model to predict the entropy for the response data of all other sounds (Methods). We observed that the prediction errors of this model were significantly lower than for a shuffled control model, implying generalized predictive ability across sound identities (Figure 4m).

Comparing the decodability of entropy with other stimulus features, such as the position of a sound in a sequence, we found a stronger representation for information content (Supplementary Figure 7f-h). Moreover, when repeating our analyses with deconvolved estimates of neural activity, we observed analogous effects, supporting their robustness (Supplementary Figure 7i-m).

These findings reveal a latent representation of stimulus identity and stimulus entropy in the space of neuronal activities evoked by our sound library. In principle, encoding of these features could either be implemented by distinct subsets of neurons in the population with selective tuning for only one of the features, or, alternatively by a more homogeneous population of neurons with mixed tuning for both features. To investigate this aspect, we characterized individual sound-responsive neurons along two dimensions: (i) their mean absolute regression weight for entropy (‘entropy weight’), reflecting their contribution to encoding information content (Methods), and (ii) their mean absolute decoding weight for discriminating sound identity (‘identity weight’), reflecting their contribution to encoding stimulus identity. Correlating entropy and identity weights across neurons revealed only a weak positive relationship (R_S_=0.039; Figure 4n), suggesting largely independent encoding of the features in the neuronal population. Overall response magnitude was more strongly associated with entropy weights than with identity weights, suggesting a stronger modulation of average response rates with information content than with stimulus identity (Figure 4o).

Together, these analyses demonstrate an abstract, stimulus-invariant representation of information content in the population activity of the auditory cortex beyond average responsiveness.

### Network mechanisms enabling the representation of information content

We wondered what neuronal mechanisms could endow a cortical network with the ability to represent the information content of an ongoing stimulus sequence. In our imaging data, the cortical representation showed two hallmarks (Figure 5a). (i) Responses adapted to repeated stimuli, i.e. the average response to a sequence of repeated sounds declined, whereas responses to variable different stimuli were maintained and even slightly increased. (ii) Information content was represented in the neuronal population geometry, with the representational axis capturing entropy and the axis capturing response magnitude being correlated but not fully aligned. Several lines of previous work point to candidate mechanisms that may enable a network to develop such signatures. On the one side, recurrent connectivity lets neural activity reflect the recent history of encountered stimuli^97^, and short-term synaptic facilitation in recurrent connections can retain recent information^98^ reshaping the effective gain of the circuit^99^. On the other side, cortical neurons show stimulus-specific adaptation, responding less to repeated than to rare inputs^47,100^, and adaptation across both afferent and recurrent synapses can reproduce stimulus-specific adaptation in recurrent models of auditory cortex^101^.

**Figure 5.**
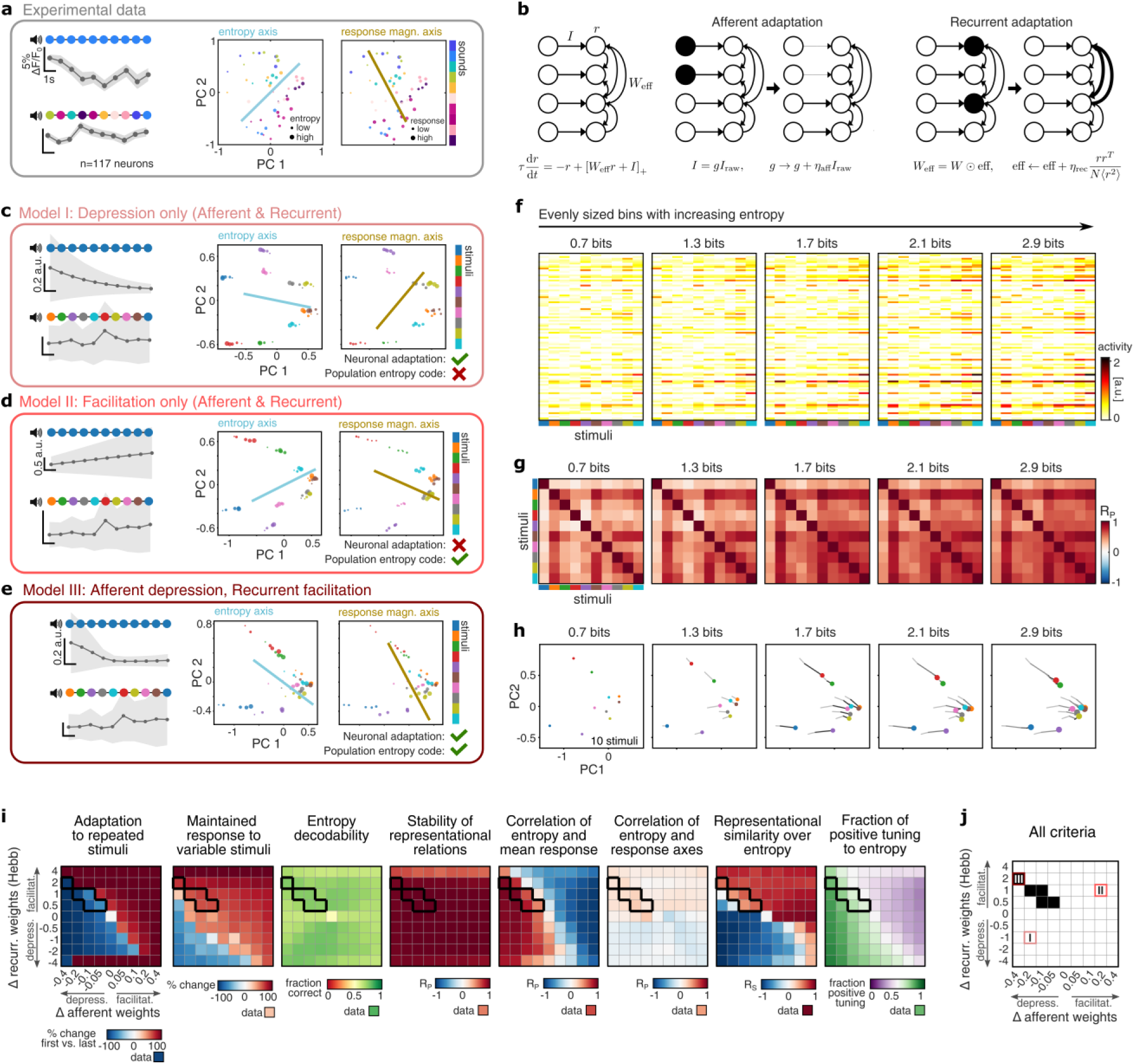
Network mechanisms enabling the representation of information content: (**a**) Experimental signatures the model aims to reproduce. Left: trial-averaged population response across a low-entropy (fully repetitive) and a high-entropy (fully variable) sound sequence. Center, right: PCA projection of the representational map, colored by stimulus identity (equivalent to Figure 4l and Supplementary Figure 7d). Marker size depicts entropy bins (center) or mean response magnitude bins (right). (**b**) Schematic of the recurrent network model with distinct mechanisms for afferent synaptic plasticity and recurrent synaptic plasticity. (**c–e**) Simulated population activity for three model settings, each displayed as in a. (**c**) Depression at both afferent and recurrent connections. (**d**) Facilitation at both afferent and recurrent connections. (**e**) Afferent depression combined with recurrent facilitation. (**f–h**) Representation of information content for the working-regime model from e: (**f**) Mean activity vectors across entropy bins. (**g**) Representational similarity matrices (i.e. correlation matrices of stimulus responses) per entropy bin. (**h**) Two-dimensional PCA projection of matrices in g (for comparison to experimental data, see Figure 4i). (**i**) Parameter scans over the two adaptation rates: the change in recurrent weights (Δ recurrent) and the change in afferent weights (Δ afferent); positive values denote facilitation, negative values depression. From left to right: adaptation to repeated stimuli, maintained responses for varying stimuli, entropy decodability, stability of representational relations, correlation of entropy and mean response, correlation of the entropy and response axes, representational similarity over entropy, and fraction of positive tuning to entropy. The working regime, fulfilling all eight criteria for the representation of information content, is outlined in black. Experimental results for all criteria are depicted in square below. (**j**) Working regime (black on white), with the parameter setting of the three example networks from c–e indicated.

Motivated by these observations, we modelled the representation of information content in a recurrent network of linearly rectified rate units, with adaptation at both the afferent and the recurrent synapses. Here, we use the term adaptation to describe changes at synapses induced by the stimulus sequence that can lead to both, depression or facilitation. The activity in the model network followed

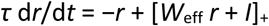

where *r* is the vector of firing rates, *τ* the time constant, [*·*]_+_ a rectified-linear activation, *I* the stimulus input, and *W*_eff_ the effective recurrent weight matrix (Figure 5b). Stimuli, sequences, and entropy were matched to those of the imaging experiment, and the network was updated after each stimulus and reset between sequences (see Methods).

In the model, adaptation could act at two complementary sites (Figure 5b). At the input, an afferent gain mechanism scaled each feed-forward input drive in proportion to its recent activation. Within the recurrent circuit, a Hebbian mechanism adjusted connections according to the co-activity of pre- and postsynaptic units. Crucially, we allowed each mechanism to change sign, so that it could act either as depression (negative values) or as facilitation (positive values) mechanism, and we systematically varied its magnitude (see Methods). With depression at the afferent and recurrent synaptic sites, we observed a marked adaptation of neural responses, declining across repeated stimuli, but no relevant population code for entropy emerged: Entropy had only a minor effect on mean activity and stimulus responses did not become more similar at higher entropy (Figure 5c). With facilitation at the afferent and recurrent synaptic sites, a pronounced population code for entropy emerged, however, adaptation of neural responses was lacking, with responses to repeated stimuli even growing (Figure 5d). Interestingly, combining afferent depression with recurrent facilitation reconciled the two effects: Responses to repeated stimuli adapted, and a population code for entropy emerged, qualitatively reproducing the experimental observations (Figure 5e).

Within this regime the model recapitulated the finer structure of the cortical representation of information content (Figure 5f–h; compare to experimental results in Figure 4). Across entropy bins, a subset of neurons became progressively more active on top of the stimulus-specific response patterns as entropy increased (Figure 5f). Representational similarity matrices retained their overall structure across entropy levels but showed broadly elevated correlations at higher entropy, indicating increased similarity between stimulus responses (Figure 5g). In a two-dimensional PCA projection, the stimulus responses retained their relative positions across entropy bins, but shifted coherently in a common direction and became more similar as entropy increased, defining a stimulus-invariant entropy axis (Figure 5h).

To map the network conditions under which this regime arises, we varied the two adaptation rates, the change in recurrent weights and the change in afferent gain, while holding network strength fixed, and evaluated the resulting representation with a set of eight metrics that we defined to quantify the representation of information content as observed in our experimental data (Figure 5i; see Methods for details): (i) The adaptation of neuronal responses across a repetitive stimulus sequence, (ii) the maintained responsiveness across a varying stimulus sequence, (iii) the overall decodability of information content from population activity, (iv) the stability of representational relations among stimuli with increasing entropy, (v) the correlation of entropy with neural mean response magnitude, (vi) the partial correlation of the representational axis for entropy and the representational axis for response magnitude, (vii) the increasing overall similarity of sound responses with entropy, and (viii) the fraction of neurons with a positive tuning to entropy. A coherent working regime, defined jointly by all metrics, emerged specifically for synaptic depression of afferent connections combined with synaptic facilitation of recurrent connections (Figure 5i,j, black outline). This working regime was robust across excitatory network configurations, but weakened as recurrent connectivity became less diverse or increasingly dominated by inhibition (Supplementary Figure 8).

Together, afferent depression and recurrent facilitation generated adaptation to repeated inputs while amplifying distributed responses to variable inputs, thus producing a stimulus-invariant population code for entropy. These findings identify the combination of afferent depression and recurrent facilitation as a minimal network mechanism for tracking the statistical structure and information content of incoming stimuli.

## Discussion

In our study, we investigated the behavioral and neuronal underpinnings of boredom in humans and mice.

First, we found that humans and mice react with avoidance behavior to monotonous sensory stimulation and that empirical entropy as a measure of information content predicts the degree of avoidance.

Avoidance behavior correlated with reported boredom in humans and was associated with low arousal and negative affect building up throughout the experiment. This is in contrast to curiosity, which is typically linked to higher arousal and positive affect^18,34,102^, and corroborates that the choice bias in our paradigm primarily reflects boredom rather than other drivers of exploratory behavior that could in principle also explain avoidance of monotony. The correlation of avoidance behavior with experienced entropy, indicates that boredom arises as an aversive cognitive response to insufficient information in sensory inputs. In our experiments, we focused on information content conveyed by the structure of the stimulus sequences while keeping other aspects of sensory information, such as the specific properties of individual sounds and additional non-auditory sensory inputs, at a low and stable level by prior habituation to the general setup. Furthermore, familiarization with the individual sounds of the sequences as well as randomization in the task structure allowed to disentangle specific responses to the information provided by the sequence structure quantified as Shannon entropy^61^. While more specialized entropy metrics may capture more detailed aspects of novelty and surprise^76^, basic Shannon entropy significantly predicted both boredom-related avoidance and neural responses, underscoring its value as a general measure of experienced information. It is important to note that the efficient transmission of information depends not only on the complexity of a transmitted sensory signal, but also on the capacity of a receiver to decode and interpret the signal^10,33,103^. We assume that basic processing and decoding of the statistical complexity in a series of inherently neutral sounds may be conserved in humans and mice, showing a low dependence on species-specific decoding abilities. Apart from observing a similar avoidance of sensory sources with low entropy in humans and mice, we found that neurophysiological measures of cortical recruitment and arousal also showed a similar correlation with information content. This further corroborates that shared neuronal mechanisms in both organisms allow to infer the complexity of its current input in order to adequately direct attention and behavior^32,34^. Our results align with previous reports in humans demonstrating a modulation of overall activity in the auditory cortex with the complexity of sound sequences^104^ together with heightened levels of arousal being important for efficient neural processing of incoming stimuli^82,105^. Furthermore, the finding that avoidance behavior in mice is driven by the experienced information content aligns with previous reports of boredom-like behaviors in various non-human animals^54,55,58,60^, including rodents^56,57,59,106^. Differences in the experiential complexity of animal boredom likely exist, nevertheless, the key feature of negative affect and additional neurophysiological signatures appear to be preserved and indicate that further mechanistic aspects of boredom may effectively be studied in animals in the future. An important implication of the above results is that the brains of both humans and mice may have the ability to sense, abstract and represent information content.

Second, we used calcium imaging of neuronal populations with single cell resolution in response to sequences with varying entropy to assess how an abstract representation of information content may emerge.

In line with the neuronal tuning of individual neurons to entropy observed in our study, previous reports have found related effects in other brain areas, identifying single neurons in the parietal cortex of monkeys^107^, or in the orbitofrontal cortex of mice^108^ that respond to the active sampling of information. Neurons showed a wide range of mixed tuning profiles to entropy and stimulus identity extending to neurons with predominant tuning to entropy. Our analyses focused on the encoding of information content emerging from the population of neurons with diverse tuning properties^109^. Observations from various brain regions support the idea that representations are formed in a lower-dimensional subspace of neuronal activity often described as a manifold^110–112^. Analyses such as representational similarity analysis (RSA) describe the structure of activity patterns within this manifold, based on the idea that the relatedness of different stimuli is encoded as a higher or lower degree of similarity between the corresponding activity patterns^92,95,113,114^. Classical studies focused on the encoding of stimulus identity and their relatedness based on perceptual or semantic properties^115–117^. Interestingly, also other, more abstract variables are mapped on the representational structure, such as sensory certainty^118^ or task contexts^119–121^. The identification of latent variables, e.g. in form of a representational axis capturing the response variability encoding a specific feature, is a common approach to make an abstract neuronal subspace functionally interpretable in form of a representational map^91,122^. Here, we identify a major representational axis, largely invariant from stimulus identity, that aligns with empirical entropy and enables the decoding of the information content from sampled sound stimuli. Such a representation allows to infer statistical variability in the structure of incoming sensory stimuli^123,124^, hence providing a basis for boredom-related behaviors. Furthermore, we found that encoding of stimulus identity along the representational axis systematically decreased with entropy, which may be related to the phenomenon of representational sharpening, describing more distinct activity patterns and enhanced discriminability for expected stimuli as compared to larger, but more stereotypical activity patterns for unexpected stimuli^125–128^. This notion supports the idea that surprising stimuli with high information content and high salience drive stronger, but less specific neural responses, facilitating subsequent learning processes at the cost of perceptual discrimination of the current sensory input^129,130^.

Third, using theoretical modeling, we identify complementary plasticity mechanisms at afferent and recurrent connections as a minimal mechanism enabling the formation of a representation of entropy in a recurrent neuronal network.

This result builds on extensive work on stimulus-specific adaptation, whereby repeated sensory inputs progressively evoke weaker responses while responses to rare or deviant stimuli are preserved^47,100^. Such adaptation has been linked to input-specific synaptic depression and can emerge from depression of feedforward inputs, recurrent connections, or both^101,131,132^. Our model extends this framework by showing that depression alone explains only one hallmark of the experimental data, where afferent depression generated adaptation to repeated inputs but did not reproduce the complexity of the observed population code for entropy. A complementary role emerged for facilitation within the recurrent circuit. Recurrent connectivity provides a natural substrate through which current population activity can depend on recent sensory history, and activity-dependent changes in recurrent efficacy can transiently retain and transform information across sequential inputs^97,98,133^. In our model, recurrent facilitation amplified the distributed activity of neurons with mixed tuning that are effectively recruited by variable stimulus sequences, thus causing population responses to shift coherently along a stimulus-invariant entropy axis while largely preserving the stimulus-specific organization. Facilitation alone, however, failed to reproduce adaptation and instead caused responses to repeated inputs to increase. Only the combination of afferent depression and recurrent facilitation reproduced both principal signatures observed experimentally, adaptation to repeated stimulation and a population representation of entropy. This division of labor in our model suggests that cortical representations of sensory statistics may emerge from interactions between plasticity mechanisms operating at distinct circuit locations. Afferent depression provides a local memory of recently encountered inputs, reducing the impact of repetitions, whereas recurrent facilitation integrates activity across changing inputs and converts their recent statistical structure into a distributed population signal. This framework complements work showing that the brain tracks sensory statistics at multiple levels, from item frequency and repetition to transition probabilities and higher-order sequence structure^134,135^ by revealing a stimulus-invariant transformation of population geometry with information content. Thus, complementary plasticity of sensory inputs and recurrent network interactions may provide a parsimonious circuit mechanism for tracking the statistical structure of ongoing sensory experience while preserving information about stimulus identity. In the future, it will be of interest to reveal the implementation of this mechanism in cortical circuits, possibly involving dynamics of synapses at shorter and longer time-scales and the emergence of local circuit motifs being disinhibited by increasing entropy in sensory inputs.

Fourth, our study together provides evidence that boredom may constitute an aversive warning signal of low information inputs to the brain.

While much of research on boredom in recent years was focused on phenomenological descriptions of its experience, it is less well understood what specific factors are inducing this aversive state. Here, we provide evidence that it is not specific physical features of sensory inputs, but rather the abstract information content conveyed by sensory stimuli that modulates boredom. Although this finding appears intuitively plausible, the application of information theoretic measures to describe the information content of sensory inputs now provides a translational framework for future studies to quantitatively describe and control the factors inducing boredom. We leveraged the mouse auditory cortex as a model to identify network mechanisms that implement the sensing and representation of information content, thereby providing a basis for future studies to elucidate how such a representation may be read out and linked to affect. Likely, multiple representations of information content for various sensory modalities may exist in parallel in the brain, and are integrated to drive boredom. For example, the insula – part of a brain network to detect the salience of sensory events^136^ – was previously shown to be implicated in boredom-related cognition^56,137^, and may constitute a neural hub integrating information content. As such, boredom can be conceptualized as a “hunger for information”, driving individuals to explore environments with varying levels of available information. In this perspective, boredom constitutes a relevant cognitive mechanism of undirected information-seeking to prevent stagnation in familiar conditions^138^, complementing directed drivers of information-seeking such as curiosity that foster the acquisition of specific information elements to fill knowledge gaps^18,22,139,140^. Thus, boredom may be interpreted as a kind of interoceptive signal driving homeostatic mechanisms ensuring a continuous stream of information to the brain which may be crucial to uphold functional representational structures at the neuronal level^141^ and to optimize the utilization of cognitive resources^8,10,142,143^. This mechanism appears particularly instrumental in respect to sensory deprivation studies reporting strong levels of distress and mental health impairment under conditions of lacking sensory stimulation^144,145^.

In summary, our study provides a characterization of the neuronal representation of sensory information content in the neocortex and its affective manifestation as boredom. Our findings connect human cognition with behavior that can be similarly observed and characterized in mice, setting a starting point for future translational studies on the neuronal and cognitive underpinnings of boredom and its role in information-seeking.

## Methods

### Human task to assess boredom-related behavior

#### Human sample for behavioral experiment

We recruited 92 healthy students from the University of Mainz for our behavioral experiment using an online recruitment system^146^. Exclusion criteria were active psychiatric or neurological disorders and insufficient German language proficiency, both assessed by self-report. The sample comprised 65 women (70.6%) and 27 men (29.4%) with an average age of 23.52 years (demographic information detailed in Supplementary Table 1). Age and gender were assessed by self-report.

#### General procedure

The human behavioral experiment was approved by the local ethics committee (Ethikkommission der Landesärztekammer Rheinland-Pfalz, processing number 2018-13164). There was no pre-registration of the study. The experiment was conducted on a single day in the facilities of the Mainz Behavioral and Experimental Laboratory (MABELLA). Written informed consent was obtained from all participants. After receiving an introduction to the study and providing consent, participants started the experiment, implemented in a custom MATLAB^®^ program which was presented on a standard computer screen. For the duration of the experiment, participants were instructed to wear on-ear headphones, through which auditory stimuli were delivered. Participants received a fixed expense allowance of €25. The study comprised the following steps:

#### Human choice task

Participants underwent a modified two-alternative choice task to assess choice behavior during sensory stimulation varying in its degree of monotony. This task has previously been shown to assess boredom-related behavior independently of other exploratory tendencies, such as curiosity^22^. Two mirror-image pushbuttons, positioned in the upper right and lower left of the screen, served as choice alternatives, and participants were instructed to make choices on these alternatives until the task ended automatically. After each choice, both buttons disappeared for 1s and a brief sound stimulus was presented through the headphones. As stimuli, we used recordings of spoken German words with neutral meaning^22^. After the stimulus interval, both buttons reappeared on their respective sides but at shifted vertical positions, so that individuals had to actively move the computer mouse to make a new decision in each trial. Participants completed three rounds of the task, each comprising 200 trials. To vary the degree of monotony at each alternative, we varied the numbers of sounds associated with each alternative, drawn from a library of 300 sounds^22^. Specifically, we tested three conditions: monotonous vs. variable stimulation (i.e. 1 sound vs. 299 sounds, mon-var), monotonous vs. monotonous stimulation (i.e. 1 sound vs. 1 sound, mon-mon), and variable vs. variable stimulation (150 sounds vs. 150 sounds, var-var). On each trial, a sound was randomly drawn from the stimulus library of the respective alternative. For each participant and round, we randomized the assignment of stimuli to the alternatives and their respective position at the left or right side of the screen to rule out side- or stimulus-specific biases. All participants first completed the mon-var condition and then completed the mon-mon and var-var condition in a randomized order. The task provided sensory stimulation without any additional reinforcement.

#### Psychometric questionnaires and ratings of boredom, affect and arousal

To assess state boredom before the task and its change over the course of the task, participants filled out the Multidimensional State Boredom Scale^16,17^ (MSBS). This questionnaire was developed to assess boredom experience in a given situation, and was previously translated into German and validated^147^. We presented the MSBS twice – before and after the choice task. Items were presented individually and in randomized order.

In addition, we used visual analog scales (VAS) to assess participants’ sentiment during the task^22,148^. Specifically, during short breaks of the choice tasks, participants were asked to rate their current degree of boredom, their current affect and their current arousal (asking “How bored/positive/aroused do you feel in this moment?”) with a slider ranging from “not at all” to “very much”. The slider had to be actively moved to enable the submission of the response. Ratings were obtained immediately before the task and at five subsequent time points, spaced approximately 30 trials apart with a random jitter of ±5 trials.

#### Psychometric ratings of sound sequences with varying information content

To assess subjective responses to stimuli varying systematically in information content, we created different sound sequences, each comprising 10 stimuli but differing in their degree of variability (60 sound sequences in total, see below for details). We then quantified empirical entropy for each of these sequences, and manually curated the sequence set to span a broad range of entropy profiles (see below). We then presented participants with all 60 sequences in a randomized order, asking them to rate each sequence with respect to perceived boredom, information content, affect and arousal (“How boring/informative/positive/arousing do you find this sound sequence?”) on a visual analog scale ranging from “not at all” to “fully”. After submitting all ratings, a 1.5s-long pause preceded the next sequence. Before sequence presentation, participants heard all ten sounds from the stimulus set three times in randomized order to familiarize them with the stimuli and their overall diversity.

### Mouse task to assess boredom-related behavior

#### Mouse sample for behavioral experiment

We used 24 male C57BL/6JRj mice obtained from Janvier laboratories for our behavioral experiments. Mice were housed in groups of four in 530 cm^2^ cages on a 12h light/dark cycle with unlimited access to dry food and water. Experiments were carried out during the light period. Mice were extensively habituated to the experimenter for 1-2 weeks before starting the experiments. At the start of the experiments, mice were approximately 6 weeks old, and the total experiment lasted ∼6-8 weeks, corresponding to an age of approximately 12-14 weeks at the end of the experiment. All animal experiments were performed in accordance with the German laboratory animal law guidelines for animal research and had been approved by the Landesuntersuchungsamt Rheinland Pfalz (approval number G 22-1-091).

#### Mouse choice task

Mice underwent a conceptually analogous choice task to assess behavior during sensory stimulation varying in the degree of monotony (see Supplementary Table 2). After several pilot experiments with reinforcement-based task paradigms, we decided to use a reinforcement-free task paradigm based on place preference. This place-preference paradigm was conceptually analogous to the human task and to previous paradigms assessing preferences for alternative forms of auditory stimulation in mice^149,150^. In our task, mice were placed in a Skinner box of 17×16.3cm (H10-11M-TC, modified by Coulbourn Instruments, Whitehall, PA, USA), separated into two mirror-image zones by a vertical wall that contained a central opening, allowing mice to switch freely between zones. The Skinner box was located in a soundproof isolation cubicle, and was illuminated by a warm-white LED. For sound delivery, we used an ASUS Xonar DX, PCI 7.1 Audio card, with an external amplifier (SLA-1, by Applied Research and Technology, TEAC Europe GmbH, TASCAM Division Wiesbaden, Germany), a modified equalizer (Applied Research and Technology, TEAC Europe GmbH, TASCAM Division Wiesbaden, Germany) and a custom-made mono loudspeaker. To assess place preference at different degrees of sensory monotony, mice underwent different task conditions that varied in the numbers of sounds associated with the two zones. A custom MATLAB® program tracked the mouse’s position during the task at 4 Hz (JIGA Stream Webcam, 1080P) and controlled sound presentation accordingly. Mice underwent the same task conditions as humans: monotonous vs. variable stimulation (i.e. 1 sound vs. 82 sounds, mon-var), monotonous vs. monotonous stimulation (i.e. 1 sound vs. 1 sound, mon-mon), and variable vs. variable stimulation (41 sounds vs. 41 sounds, var-var). The overall stimulus library consisted of 83 different sounds comprising 70ms-long pure tones and complex sounds, 1s-long environmental sounds, and several 1s-long recordings of spoken words, all presented at a sound pressure level of 65dB. Sound onsets were separated by an inter-stimulus interval of 3 seconds. In addition to the above task conditions, we included silence as a negative control for auditory stimulation, and loud white noise stimulation (1s-long, 75dB) for which we expected avoidance behavior^69,70^. This led to the additional task conditions: silence vs. silence (sil-sil), white noise vs. silence (noise-sil), white noise vs. variable (noise vs. 83 sounds, noise-var) and silence vs. variable (silence vs. 83 sounds, sil-var) (see Supplementary Figure 2). Mice subsequently underwent multiple 40min-long sessions under different task conditions. For each session, the sound assignment and the assignment of conditions to the right or left zone in the box was randomized. At the beginning of the experiment mice underwent all seven task conditions (mon-var, mon-mon, var-var, sil-sil, noise-sil, noise-var, sil-var) in pseudorandomized order until they completed 6 sessions in each condition. Subsequently, mice were exposed to 12 additional sessions of the mon-var condition. Each mouse completed two sessions per day. Note that for the noise-sil condition, we tested only a smaller subset of 16 mice.

### Quantification of entropy in the choice task

To quantify the amount of experienced sensory information in the task, we quantified the diversity of stimuli sampled by participants or mice up to each point in the task. For the human data, we considered the sounds sampled across all trials in a given task condition. For the mouse data, we split the session data into discrete time bins before each sound presentation (every 3 seconds), treating this binned data equivalently as trials for our analyses. We then computed Shannon’s empirical entropy as a measure of information content^61^, based on the relative frequencies of sounds sampled from each alternative. Specifically, we computed the entropy *H* from alternative *k* at a given trial *t* as 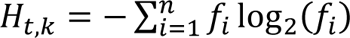, where *f_i_* denotes the relative frequency of sound *i* among all sounds sampled from alternative *k* up to trial *t*. *n* denotes the number of all unique sounds sampled from alternative *k* up to the current trial (Supplementary Figure 1e). Thus, repeated sampling of the same stimulus generally decreased entropy, whereas sampling of novel stimuli increased entropy.

### Creating sound sequences with varying information content

To assess the neural representation of information content in the cortex of humans and mice, we used empirical entropy^61^ to create sequences of sounds that systematically varied in their complexity. Specifically, we used an initial library of ten sounds and generated sound sequences by sampling from this library with replacement. For the human experiment the initial sound library consisted of ten 500ms-long complex snippets of naturalistic environmental sounds or snippets of animal vocalizations with qualitatively distinct acoustic characteristics. The initial mouse library consisted of ten 70ms-long sounds (5 pure tones and 5 complex sounds) that were previously shown to evoke distinct activity patterns in the auditory cortex^90,151^. From the initial library, we created a set of 100 sound sequences with a length of ten stimuli and intervals of 1 second between sound onsets. We then computed the empirical entropy for each sound sequence^61^ based on the frequency of each sound relative to all stimuli presented up to that sequence position. Here, we assumed initial silence before the sequence onset as a ‘blank’ stimulus, and computed empirical entropy *H* for each sound *s* as 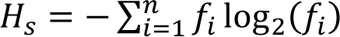, where *i* represents each of the *n* unique stimuli sampled until position *s* in the sequence, and *f_i_* represents the relative frequency of each unique sampled stimulus (see Supplementary Figure 3). This procedure provided a metric of entropy that generally decreased when a stimulus was presented repetitively, and increased when new stimuli were presented. Moreover, entropy generally tended to increase with the number of sounds presented in a sequence. The highest entropy values occurred for sequences containing many distinct sounds sampled independently with approximately uniform probabilities. Based on these entropy profiles, we manually selected and refined 20 sequences spanning a broad range of overall entropy values and within-sequence entropy dynamics. To avoid stimulus-specific response biases, we then shuffled the assignment of sound identities to the sequence patterns, creating three and five different versions of the 20 sound sequences for the human and mouse experiment, respectively (see Supplementary Figure 4).

### Human electroencephalography and pupillometry with sound sequences with varying entropy

#### Human sample for EEG and pupillometry experiment

We recruited 12 healthy students from the University of Mainz for the combined EEG and pupillometry experiment through written announcements. Exclusion criteria were active psychiatric or neurological disorders as well as insufficient German language skills, both assessed by self-report. Participants reported no use of psychotropic medication. The sample comprised 8 women (66.7%) and 4 men (33.3%), with an average age of 24.9 years (demographic information detailed in Supplementary Table 3). Gender and age information was collected through self-reports.

#### General procedure and presentation of sound sequences

The experiment was approved by the local ethics committee (Ethikkommission der Landesärztekammer Rheinland-Pfalz, processing number 2018-13164_8). There was no pre-registration of the study. The experiment was conducted on a single day in the facilities of the Neuroimaging Center Mainz. Written informed consent was obtained from all participants. After receiving an introduction to the study and providing consent, participants began the experiment, which was implemented in a custom MATLAB® program and presented on a standard computer screen, approximately 80 cm from the participants’ head. Participants received a fixed expense allowance of €25. The study comprised the following steps: First, participants completed different psychometric questionnaires assessing general information, personality traits^147,152–156^, and state boredom^16,147^. While participants filled out the questionnaires, we prepared the 64-channel EEG and pupillometry recordings (see below for details). After testing the EEG electrodes for sufficiently low impedance (<10 kΩ) and calibrating the pupillometer, we started the recording and began to present sound sequences with varying entropy. Per participant, 60 sound sequences (20 different sound sequence patterns in 3 versions, see above and Supplementary Figure 4a) were presented in randomized order. Each sequence consisted of 10 sounds, i.e. 10 second duration, and was followed by an inter-sequence interval of 20 seconds. Sounds were delivered via two external loudspeakers located in front of the screen, approximately 95cm away from a participant’s head. During the experiment, participants were presented with a blank screen showing a central black fixation cross and were instructed to maintain fixation and minimize movement. After the auditory sequence presentation, participants provided subjective ratings of all sound sequences in line with the above experiment in the larger sample (see above). In addition, participants underwent an equivalent presentation of visual stimulus sequences (using neutral images of everyday objects^157,158^) and two rounds of a choice task between monotonous and variable stimuli in auditory and visual modality, while we recorded EEG and pupil size. Before and after the choice task, participants again rated state boredom on the MSBS. In the present study, we restricted our analysis to the auditory sequence presentations, enabling direct comparison with the mouse experiments described below.

#### Electroencephalography (EEG) and pupillometry

EEG data were acquired using a 64-channel cap with passive Ag/AgCl electrodes (EasyCap, Germany) positioned according to the extended international 10–20 system^159^. Signals were amplified using a 24-bit NeurOne Tesla amplifier (Bittium, Finland), powered by a 7.2 V battery, and sampled at 1000 Hz. During acquisition, FCz served as the online reference and POz as the ground electrode. Electrode impedances were maintained below 10 kΩ before recording.

Simultaneously, pupil diameter was recorded using a head-mounted eye-tracking system (Pupil Labs Core, Pupil Labs) at a sampling rate of 30 Hz. A standard calibration procedure was performed at the beginning of each session. Recordings were conducted under ambient room lighting conditions. EEG and pupillometry data streams were synchronized via shared event markers to ensure precise temporal alignment.

### Preprocessing of human imaging data and quantification of evoked responses

#### Electroencephalography data

Preprocessing was performed using custom MATLAB R2023a scripts utilizing EEGLAB^160^ and FieldTrip^161^ functions. Continuous recordings were downsampled to 250 Hz to optimize computational efficiency while maintaining temporal resolution for ERP analyses. Line noise (50 Hz and its harmonics) was removed using Zapline-plus^162^, an adaptive extension of the Zapline algorithm^163^ for automated, parameter-free spatial filtering of power-line artifacts. To improve independent component analysis (ICA) stability, the data were high-pass filtered at 1 Hz using a two-pass zero-phase FIR filter (pop_eegfiltnew, order 1650). Data were re-referenced to a full-rank common average reference^164^ to preserve rank for subsequent spatial decompositions. EEG data were subsequently processed using the Generalized Eigenvalue De-Artifacting Instrument (GEDAI), a model-based denoising approach that uses a theoretical leadfield matrix to constrain signal decomposition^165^. Assuming that neural activity is consistent with a biophysically plausible forward model of head volume conduction, GEDAI applies generalized eigenvalue decomposition to isolate components that fit the model and attenuate non-neural activity. This method was applied prior to ICA to maximize decomposition stability. ICA was performed on the continuous data using the extended Infomax (runica) algorithm, with PCA dimensionality reduction set equal to the effective rank of the data^164^. Independent components were classified using ICLabel^166^ and rejected if artifact probabilities met conservative thresholds (muscle, heart, or channel noise ≥ 0.97; eye ≥ 0.90), or if the probability of brain activity was ≤ 0.01 (number of rejected components per participant: mean=3.75 ± SD=1.30). All flagged components were visually inspected to verify their classification as non-physiological. Following ICA correction, we visually inspected channel-wise power spectral density (4s segments, 80% overlap, Hanning window) plots for each participant to confirm signal quality. Finally, continuous data were segmented into epochs from −250 to 1000ms relative to stimulus onset and smoothed with a moving mean filter over 5 samples (i.e. 20ms). Because artifact attenuation was performed at the continuous-data level using GEDAI and ICA, no additional epoch-level rejection procedures were applied.

To quantify auditory evoked cortical potentials, we considered the average evoked potential over the channels Fz and Cz. As a measure of sound-evoked activity, we quantified the mean P200 EEG potential of these electrodes in the 100-250ms period after the onset of each stimulus, averaging across all versions of the sequences. This procedure quantified components of the auditory evoked potentials that have previously been associated with sensory expectation and surprise signals^49,50,83^. For comparison, we analogously computed the N100 signal in a window of 80-130ms after sound onset, and the P300 signal in a window of 250-350ms. In analogy to the above analyses, we also tested the correlations of N100 and P300 with entropy, but observed only weaker modulations (see Supplementary Figure 5b,c). Therefore, we decided to use the P200 component as the main readout of auditory evoked EEG activity in our study. As a measure for the overall neural response to a given stimulus sequence, we also computed the mean P200 signal over the ten sound presentations in each sequence.

#### Pupil data

For the preprocessing of the human pupil data, PupilPlayer (version 3.5.1; Pupil Labs) was used to extract absolute pupil-size time series from each recording. For each sample in the time series, we also obtained a confidence index provided by the analysis, ranging between 0 and 1, indicating the quality of the ellipse fit for each frame. We excluded participants for whom ≥40% of samples had confidence values <0.7, leaving eight participants for analysis (see Supplementary Table 3 for details). For each individual’s data, we excluded outlying single samples more than 3 SD from the overall median pupil size, and interpolated the excluded data linearly. To quantify pupil responses for all sound presentations across sequences, we first smoothed the pre-processed time series data with a moving mean filter over 10 frames (i.e. 333ms), and then subtracted the mean baseline pupil size in the 1 second interval before each sequence onset from the time series of the respective sequence. We then computed the mean pupil response for each sound as the mean pupil size in the 1 second period after the onset of each sound in the sequence, averaging across all versions of the sequences. As a measure for the overall pupil response to a given sound sequence, we also computed the mean over the ten individual sound-evoked pupil responses.

### Mouse imaging experiments with sound sequences of varying entropy

#### Mouse sample

We used 5 male C57BL/6JRj mice obtained from Janvier laboratories for our imaging experiments. After the surgeries, mice were single-housed in 530 cm^2^ cages on a 12h light/dark cycle with unlimited access to dry food and water. Experiments were carried out during the light period. Mice were habituated to the experimenter for one week before starting the experiments. At the start of the surgeries, mice were approximately 6 weeks old, and the imaging experiments were started 3-4 weeks later, lasting for approximately 4 additional weeks. All animal experiments were performed in accordance with the German laboratory animal law guidelines for animal research and had been approved by the Landesuntersuchungsamt Rheinland-Pfalz (approval number G 22-1-091).

#### Stereotaxic injections and cranial window implantation

We followed the experimental procedures described in detail previously^90,151^. Briefly, mice were anesthetized with isoflurane (maintained at 1.2-1.5% at a flow rate of around 200mL/min) using a vaporizer (High Precision Instruments, MT; Univentor 400 Anesthesia Unit), and stereotaxic injections were performed perpendicular to the surface of the skull. Virus solution consisted of a mixture of two different rAAVs (rAAV8-hSyn-GCaMP6m-WPRE-hGHpolyA; titer: 1.35 × 10^11^ viral genomes (VG)/ml; rAAV8-hSyn-H2BmCherry-hGHpolyA; titer: 2 × 10^13^ VG/ml) in PBS. The viral mixture was loaded into a glass pipette and injected at five locations (170 nL per site at 20 nL/min; 850 nL total volume) along the anterior–posterior axis with coordinates: 4.4, −2.5/−2.75/−3.0/−3.25/−3.5, 2.5 (in mm, caudal, lateral, and ventral to Bregma) to cover the right auditory cortex. Two weeks after the injection, mice were anesthetized using isoflurane (maintained at 1.2-1.5 %) and mounted on a stereotaxic frame with a custom-made v-shaped head holder. After the parietal and temporal bone were exposed, the skull above the auditory cortex (2×3 mm) was gently drilled and the bone was carefully lifted. The craniotomy was covered with a small coverslip secured with dental cement. To position the window plane perpendicular to the objective under the microscope, a custom-made titanium head post was mounted on the frontal bone and embedded with dental cement. After the surgical procedure, mice recovered for at least one week before further handling.

#### Habituation and sound presentation

Before starting the neuronal imaging, we habituated mice to head fixation and sound presentation for approximately one week. We gradually increased the time for which mice were head-fixed while presenting them with repetitions of the ten sound stimuli used to create the sound sequences. Sounds were presented in a random order, at a sound pressure level of 65dB and with inter-stimulus intervals of 1.5 seconds. Sound sequences were delivered using the system described in detail in previous studies from our laboratory^90,151^. In short, all sounds were delivered in a free field setting through a ribbon loudspeaker positioned 25 cm from the mouse’s head in a soundproof booth. We presented mice with the same 20 sound sequence patterns that we used in the human experiment, assembled from a library of 10 short sounds (70ms duration, 5 pure tones, 5 complex sounds) which were previously shown to elicit distinct activity patterns in the auditory cortex^90,151^. For the mouse experiment, we created 5 different versions of the 20 sound sequences, each containing intervals of 1 second between the onset of subsequent sounds, and inter-sequence intervals of 20 seconds (see Supplementary Figure 4b).

#### Mesoscopic calcium imaging, pupillometry and movement assessment

We followed an established protocol for widefield calcium imaging in the auditory cortex^88^. The mesoscopic optical imaging setup comprised a CCD camera (Vosskuehler, Germany, CCD1200QD), attached to a macroscope consisting of two objectives placed face-to-face (Nikon 135 and 50 mm), and a blue excitation LED (470 nm). Mice were placed under the camera and fixed via their head post. We recorded mesoscopic calcium activity from the right auditory cortex of awake, passively listening mice at 10Hz and 200μm below the cortical surface. GCaMP6m was excited with a 470nm LED while sound sequences of varying entropy were presented Each of the 100 sequence-and-version combinations was presented five times, where the order of sequences within each repetition was randomized, corresponding to a total number of 5000 sound presentations (10 sounds per sequence, 20 sequences, 5 versions, 5 repetitions). Simultaneously, pupil size was recorded using a camera-based eye-tracking system (Raspberry Pi Camera Module 3 NoIR, 12MP, RPI-CAM3-NO, sampling rate of 5 Hz) positioned laterally to capture the eye ipsilateral to the imaged hemisphere. The eye was shielded from the blue excitation light with a curved piece of aluminum foil. In addition, body movement was coarsely monitored using a distance sensor (Waveshare, Infrared Reflective Sensor, LM393, sampling rate of 9 Hz) positioned behind the animal to detect movements of the back and torso. All data streams (calcium imaging, pupil size, and movement) were synchronized via shared triggers and timestamps to ensure precise temporal alignment. Recordings were conducted in a dark soundproof box that was only illuminated by the blue LED used for mesoscopic imaging. One mouse was excluded from the mesoscopic calcium and pupil analyses because of excessive movement, poor image quality, and substantial occlusion of the pupil by the eyelid (see Supplementary Figure 4).

#### Intrinsic imaging

We used intrinsic signal imaging to map auditory fields and guide the selection of fields of view (FOVs) for two-photon calcium imaging. We followed a previously reported protocol from our laboratory^90,167^: Mesoscopic optical imaging was conducted using a CCD camera (Vosskuehler, CCD1200QD; frame rate of 25Hz) with macroscope objectives. Intrinsic signals were recorded at 200μm below the brain surface under illumination with near-infrared LEDs (780nm) under isoflurane anesthesia (concentration kept at 1-1.2%), while presenting trains of 18 pure tone pips (80ms-long individual pips with 20ms smooth gaps; 1.8 s of the total stimulus duration) with different pure tone frequencies (1, 2, 4, 8, 16, 32 and 64 kHz, respectively) and white noise bursts in 30 randomized trials. The sound intensity was set at a sound pressure level of 70dB. The change in light reflectance between pre- and post-stimulus frames was computed and averaged across trials (for an exemplary tonotopic map see Supplementary Figure 6a, for the distribution of auditory fields across FOV see Supplementary Figure 6b).

#### Two-photon calcium imaging

After completing the mesoscopic imaging, two-photon calcium imaging in the auditory cortex of awake head-fixed mice was performed. The imaging configuration and procedure have been described in detail previously^90,151^. Briefly, two-photon calcium imaging was performed with a commercial microscope (Ultima IV, Bruker Corporation) with a 20×-objective (NA=0.95, Olympus) and a tunable pulsed laser (Chameleon Ultra, Coherent) in a soundproof chamber. Both GCaMP6m and mCherry were co-excited at 940 nm. Time-series imaging was performed using a field of view (FOV) of 367×367μm at a frame rate of 5Hz. Across multiple days, with imaging sessions typically separated by approximately two days, 6-11 different FOVs were imaged per mouse in cortical layer 2/3 (about 125-175μm depth from the cortical surface). For a given FOV, all 5 versions of the 20 sound sequences were presented, corresponding to a total number of 1000 sound presentations. Details on the structure of our dataset and the FOV distribution across mice are presented in Supplementary Table 6.

### Preprocessing of mouse imaging data and quantification of evoked responses

#### Mesoscopic calcium data

The mesoscopic image recordings were first corrected for movement artifacts by aligning them to a reference frame at the beginning of the recording using rigid translational image registration, and smoothing the time series across time using a moving mean over 4 frames (i.e. 200ms). Then, we computed ΔF/F_0_ traces for all pixels in the image, defining F_0_ as the mean signal intensity in the 5 seconds before the first sound onset of a sequence. For each individual field of view, we then identified sound-responsive pixels by taking the mean evoked response over all frames as a template and defining a mask manually to cover the most responsive regions. We verified that this mask matched the evoked response patterns obtained by intrinsic imaging in the corresponding mouse. Then, we averaged all pixels within the mask across the full time series, providing a measure of globally evoked responses in the auditory cortex over time. To compute the evoked calcium response to each individual sound, we averaged the ΔF/F_0_ signal in a 600ms time period after sound onset. Averaging the evoked response over all sounds of a given sequence provided a metric for overall activity in response to a sequence with a defined entropy profile.

#### Pupil size data

From the raw pupil recordings, we extracted the pupil size (diameter) over time using a published analysis pipeline based on MATLAB^168^. Briefly, this algorithm performs threshold-based segmentation to isolate the pupil region, followed by contour detection to identify its boundary. The detected pupil shape was approximated via ellipse fitting, providing an estimate of pupil size over time. The time-series data were smoothed using a moving mean filter over an interval of 500ms. In addition, for the time series data of each sequence, we subtracted the baseline pupil size, computed as the mean over an interval of 5 seconds before sequence onset. To compute the evoked pupil response to a given sound, we then calculated the mean over a 600ms window after each sound onset.

#### Movement data

To obtain a metric of evoked movement from the raw distance recordings, we first computed the derivative of distance values and then removed individual outlying samples that showed extreme values (i.e. due to a movement of the tail towards the distance sensor). We also smoothed the data over time using a moving mean over an interval of 500ms. To correct our movement estimates for individual differences in sensor or body position, we computed for each mouse the z-score of the change in measured distances during each sound sequence and the preceding 20 seconds baseline interval. We then used the absolute value of this z-score, providing a metric of the overall movement of a given mouse over the course of presenting the sound sequence, relative to its current activity state. To quantify sound-evoked movement, we computed the mean absolute z-score in the 600ms after each sound onset.

#### Two-photon calcium data

Before applying an individual cell tracking algorithm, global xy-image displacement induced by movement was corrected by a cross-correlation based method^169^. Regions of interest (ROIs) were semi-automatically selected by a custom-written MATLAB program, and were manually corrected by a human expert. ROIs usually comprised a set of several hundred points marking the centers of neuronal nuclei. To track individual ROIs across frames, we followed a standardized approach described in a previous study from our lab^151^. A two-dimensional affine transformation was applied to register ROIs across frames within each FOV. To include only cells with a reliable nucleus signal (H2B:mCherry channel) in the analysis, we applied four quality criteria described in our previous study^151^. To correct fluctuating background fluorescence from somatic Ca^2+^ signal, the out-of-focus neuropil signal was calculated as the average fluorescence value of an area surrounding each individual region of interest and subtracted from the time series after multiplication with a contamination ratio as described before^170^. The contamination ratio was calculated independently for each imaging plane in the experiment. To calculate ΔF/F_0_, F_0_ was defined as a moving rank-order filter, namely the 30th percentile of the 200 surrounding frames (100 before and 100 after, i.e. 20 seconds before and after).

### Network model of information content

#### Network model

We modelled a recurrent network of *N* = 100 rate units. The activity dynamics followed

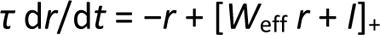

where *r* is the vector of firing rates, [*·*]_+_ denotes half-wave rectification (i.e. rectified-linear activation), *I* is the stimulus input, and *W*_eff_ is the effective recurrent weight matrix. The time constant was *τ* = 1 and the equations were integrated using Euler’s method with step size *dt* = 0.01 for 1000 steps per stimulus, which was sufficient for the activity to reach a steady state.

The recurrent weight matrix was initialized as a Gaussian random matrix with a specified entry-wise mean and standard deviation, which were then held fixed throughout the simulation. Entries were drawn as

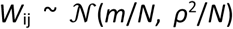

with *m* = 0.25 and *ρ* = 0.9, so that *m* sets the mean (i.e. common-mode) eigenvalue of the matrix and *ρ* the spectral radius of its random part. After the initial draw, and again after every within-sequence weight update (see below), the matrix was renormalized to restore exactly this mean and standard deviation,

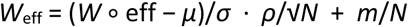

where *μ* and *σ* are the empirical entry-mean and entry-standard-deviation of *W* ∘ eff. Ten stimuli (A-J) were constructed as sparse input vectors: for each stimulus a distinct, randomly chosen 10% of units (10 of 100) was assigned an input magnitude drawn independently from *U*[0, 1], with all remaining units receiving zero input. Stimulus identities were fixed for a given network throughout the simulation. The network was presented with stimulus sequences identical in structure to those used in the mouse imaging experiment: the same 100 sequences (5 sequence versions × 20 sequence patterns) of length 10, preserving the exact temporal statistics, with the synthetic sparse vectors above substituted for the experimental stimuli. Contextual entropy at each sequence position was defined exactly as in the experimental analysis. After each complete sequence the adaptation state was reset (both adaptation variables, below, returned to baseline) while the base weight matrix *W* remained fixed.

#### Adaptation mechanisms

During sequence presentation the network adapted through two concurrent mechanisms. After each stimulus, the corresponding adaptation variable recovered exponentially toward its baseline with a time constant of *τ*_rec_ = *τ*_aff_ = 10 sequence steps.

*Recurrent Hebbian plasticity (on the outputs):* A matrix of multiplicative synaptic efficacies, eff (initialized to the all-ones matrix), was updated by a normalized outer-product rule on the response,

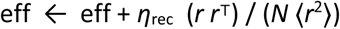

followed by exponential recovery toward unity,

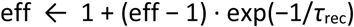

The effective recurrent matrix was then *W*_eff_ = renorm(*W* ∘ eff) using the renormalization above, so that plasticity reshaped the connectivity *structure* while the mean eigenvalue and spectral radius were held fixed.

*Afferent gain plasticity (on the inputs):* A per-input gain vector *g* (initialized to ones) scaled each feed-forward synapse in proportion to the raw input magnitude it received,

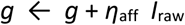

with exponential recovery toward unity,

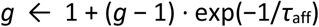

The effective drive entering the dynamics was then *I* = *g* ⊙ *I*_raw_. Stimulus dependence therefore emerged from the input-magnitude pattern itself.

#### Parameter sweeps

We performed a two-dimensional sweep over the two adaptation rates, forming a 9 × 9 grid: the recurrent Hebbian rate *η*_rec_ ∈ {−4, −2, −1, −0.5, 0, 0.5, 1, 2, 4} (Δ recurrent weights) and the afferent gain rate *η*_aff_ ∈ {−0.4, −0.2, −0.1, −0.05, 0, 0.05, 0.1, 0.2, 0.4} (Δ afferent weights). For both axes, positive values correspond to facilitation and negative values to depression. All other parameters were held fixed at the values given above. Metrics were averaged across 100 independently initialized networks per grid condition.

#### Metrics quantifying the observed and modelled representation of information content

We computed eight metrics characterizing how population activity related to contextual entropy. For metrics that required entropy binning, responses and entropy values were assigned to one of five equally spaced bins spanning the observed entropy range.

#### Adaptation to repeated stimuli

To quantify the response dynamics to maximally redundant stimulus sequences, we computed the signed percentage change in mean population activity (last versus first stimulation) for the minimum-entropy, fully repetitive sequence, averaged over its five versions. Negative values indicate adaptation.

#### Maintained responses to varying stimuli

To quantify the response dynamics to maximally diverse stimulus sequences, we computed the same signed percentage change in mean population activity for the maximum-entropy sequence (all different stimuli), averaged over its five versions. Positive values indicate facilitation.

#### Entropy decodability

We assessed whether network activity carried decodable information about momentary entropy by training a linear discriminant classifier to separate low-from high-entropy trials (median split) following dimensionality reduction to 20 principal components; accuracy was estimated by 5-fold cross-validation.

#### Stability of representational relations

For each entropy bin, a representational similarity matrix across the 10 stimuli was computed as the pairwise Pearson correlations between mean stimulus responses; stability was the mean Pearson correlation between all pairs of bin-level similarity matrices.

#### Correlation of entropy and mean response

The Pearson correlation between mean population activity and mean entropy, computed across entropy bins.

#### Correlation of entropy and response axis

Ridge regression was used to estimate the population axis that best predicted entropy and, separately, the axis that best predicted mean activity; the metric was the Pearson correlation between the two resulting weight vectors.

#### Representational similarity over entropy

The mean within-bin representational similarity (mean of the upper triangle of the bin similarity matrix) was computed per bin, and its Spearman rank correlation with mean bin entropy was reported.

#### Fraction of positive tuning to entropy

The fraction of units whose activity was positively correlated with entropy across trials, assessed via a per-unit Spearman rank correlation.

#### Working regime

A grid condition was classified as lying within the working regime when the network reproduced the qualitative signatures observed in the experiment. We therefore required all metrics to point in the expected direction: activity adapted across the fully repeated (minimum-entropy) sequence (change ∈ [−100, 0]%); activity facilitated across the maximally changing (maximum-entropy) sequence (change ∈ [0, 100]%); entropy was decodable above chance (LDA accuracy > 0.5); representational relations were stable across entropy levels (mean inter-bin RSM correlation ≥ 0.5); mean response correlated positively with entropy (r ∈ [0, 1]); the entropy-encoding and mean-activity axes were positively correlated (r ∈ [0, 1]); representational similarity increased with entropy (Spearman ρ ≥ 0); and a majority of units were positively tuned to entropy (fraction > 0.5). Because our aim was a mechanistic understanding of how these signatures arise rather than a precise reproduction of the experimental data, we sought this qualitative correspondence rather than a quantitative fit. Within a regime meeting the above criteria, network activity is reliably modulated by both stimulus identity and contextual entropy, with the two signals occupying partially overlapping but distinguishable subspaces of population activity.

### Statistical analyses

All analyses were conducted using the MATLAB^®^ statistics and machine learning toolbox (The Mathworks Inc., Natick, Massachusetts, USA, version R2022a). Full details of the statistical tests, sample sizes, and p-values for all analyses are provided in the separate Statistical Information file.

### Quantification of preference and other parameters in the choice task

From the behavioral data of humans and mice, we quantified the choice preference by computing the mean probability of choosing a fixed reference alternative in each respective task condition over a moving bin of 10 trials (human) or 1 minute (mouse). The reference alternative depended on the task condition (e.g. in mon-var, the variable alternative was chosen as a reference), whereas for conditions with equivalent stimulation on both sides, we randomly assigned the right or left alternative as reference for each mouse individually. For mice, we further computed the mean distance traveled as well as the frequency of switching between zones in an equivalent time bin of 1 minute. To compare the choice preference across task conditions at a steady state level, we quantified the mean preference of participants/mice in the second half of the tasks during the last two sessions of each condition.

### Self-report questionnaires and psychometric ratings

To compare state boredom in humans across the experiment, we calculated the total score of the Multidimensional State Boredom Scale^16^, as well as scores in all the MSBS subscales (Disengagement, Inattention, Low Arousal, High Arousal, Time Perception), and compared these metrics between the pre-and post-task assessment. To investigate sentiment over the course of the choice task, we compared the raw visual analog ratings of boredom, affect, and arousal, all of which ranged between a minimum value of 0 and a maximum value of 1.

### Quantifying the relationship between experienced entropy difference and choice probability

We discretized the mouse preference data, segmenting the time series into 3-s bins aligned to sound onsets. For each time bin, we then computed the chosen alternative as the zone in which the mouse spent more time in the current time bin. This provided a trial-based data structure comparable to the human choice task data. For each trial, we then quantified experienced entropy for both alternatives, based on the variability of previously sampled stimuli (see above). Note that despite a large degree of qualitative congruence, the absolute values for sampled entropy varied between humans and mice due to differences in the size of stimulus libraries in the task and a higher tendency of mice to stay within a given zone independent of sensory stimulation. We then calculated the difference in entropy between alternatives for each trial. For each trial *t*, we also computed the choice probability in the next trial *t+1* for the higher-entropy alternative. Then, we sorted the single trial data of each individual into bins of increasing entropy difference (edges human: [0.2, 0.85, 1.5, 2.15, 2.8, 3.45, 4.1, 4.75, 5.4, 6.05] bits, edges mouse: [0.65, 1.3, 1.95, 2.6, 3.25, 3.9, 4.55, 5.2] bits), and computed the mean sampled entropy difference and the corresponding choice probability for each individual.

### Logistic regression model to quantify the effect of experienced information content on choice behavior

We used the trial-segmented data of humans and mice to quantify the impact of experienced information content, i.e. entropy, on individual choice behavior in the mon-var condition and to compare this effect with other typical decision factors. A similar logistic regression model has previously been shown to successfully capture behavior in an equivalent choice task^22^. Specifically, we applied a logistic regression model, expressing the choice probability for alternative *a* on trial *t* as 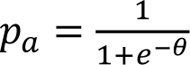, where *θ* = *w*_1_ + *w*_2_Δ*H* + *w*_3_*C_a_*. In this equation, θ expresses a linear combination of different predictors and their corresponding weights that independently influence choice behavior. *Bias*: *w*_1_ represents a stimulus-independent idiosyncratic bias towards a given side^64,65^. *Entropy difference*: *w*_2_Δ*H* expresses the sensitivity of an individual to adjust its choice to differences in experienced entropy (Δ*H*). *Inertia*: *w*_3_*C_a_* represents a stimulus-independent tendency to repeat the previous choice^66–68^, where *C_a_* takes either a value of +1 if the choice in the previous trial *t-1* was in favor of alternative *a*, or a value of −1 if the previous choice was not in favor of alternative *a*. For each participant/mouse, we fitted the weights of all parameters in the regression over all trials in the mon-var condition, using a maximum likelihood approach. As a metric of model goodness, we then evaluated the fraction of correctly predicted trials from the fitted regression model for each participant. To compare the effects of the different regressors, we estimated the contribution of each parameter to individual choice behavior by considering the weights scaled by the respective parameter values. In particular, we computed the contribution of idiosyncratic choice bias as *contribution_bias_* = |*w*_1_|, the contribution of experienced entropy as *contribution_entropy_* = |*w*_2_*ΔH*|, and the contribution of previous choice on the current decision as *contribution_inertia_* = |*w*_3_*C_a_*|. For the entropy term, we computed |*w*_2_Δ*H*| on each trial and averaged these values across trials.

### Sentiment of sound sequences with low versus high information content

For the sentiment of sound sequences with varying information content, we z-scored ratings of each feature (boredom, information content, affect, and arousal) across the 60 sequences for each participant. This procedure adjusted sentiment estimates for individual response biases, such as systematic differences in scale use across participants. We then compared the mean ratings of each participant across the five highest-entropy sequences with those of the five lowest-entropy sequences.

### Correlation of cortical responses, arousal, and the mean information content of sound sequences

To assess how strongly overall auditory evoked cortical responses and pupil response represent sensory information content in humans, we quantified the mean evoked P200 potential across the Fz and Cz electrodes, as well as the mean evoked pupil response across participants to all 20 sequence patterns (see above). We then correlated these measures with the mean entropy of the sequence patterns (Pearson correlation). For the mouse data, we conducted an equivalent analysis for the mean evoked mesoscopic calcium response, the mean pupil response and the mean movement evoked by all 20 sequence patterns.

### Correlation of cortical responses, arousal, and information content of single stimuli

To investigate how evoked cortical activity, pupil response and movement reflect entropy of single short sensory events, we conducted a multiple linear regression (MATLAB function *fitlm*), modeling the entropy of all single sound presentations – pooled across all sequences, versions and participants – as a linear combination of the evoked EEG responses and pupil responses (human), or as a linear combination of evoked mesoscopic calcium responses, pupil responses and movement (mice), respectively. We conducted the regression including an intercept, and included only individuals that provided data for all regressors (see Supplementary Table 4 for statistical details).

### Sound responsiveness of single neurons

To classify the sound responsiveness of single neurons with good imaging quality (n=7,392), we used the raw ΔF/F_0_ traces, pooled across all sequence presentations, and extracted all frames corresponding to the presentation of a specific sound and the 600ms interval after sound onset. We then compared the calcium signal of each cell for these sound-related frames (stimulus-related activity) to the calcium signal across all other frames outside of any stimulus presentations (stimulus-unrelated activity) using a Wilcoxon rank sum test. A neuron was classified as significantly *excitatory* or *suppressive* to this specific sound, if the median stimulus-related activity was higher or lower than the median stimulus-unrelated activity and if the Benjamini-Hochberg-corrected p-value was <0.05 after correcting for multiple comparisons against the number of all cells and sounds. We repeated this procedure for all other sounds in the sequence library (n=10), classifying neurons as having (i) no significant response, (ii) an excitatory response, or (iii) a suppressive response^89,91^. In total, n=5,454 neurons showed a significant response to at least one sound. For our further analyses, we focused on these responsive neurons, omitting the data of neurons without significant response. Details on the average response patterns across all sounds and the distribution of response types across neurons are summarized in Supplementary Figure 6c-g.

### Deconvolution of two-photon calcium data

Even though we conducted our main analysis with the raw ΔF/F_0_ traces, we also replicated our key findings using deconvolved activity estimates. We used the algorithm published by Vogelstein and colleagues for deconvolution^171^. This algorithm, as is typical of classical deconvolution methods, reduces decreases in calcium-dependent fluorescence to values of zero, therefore often leading to a loss of suppressive responses in the data. To counteract this effect, we computed a deconvolved response to each sound by averaging activity over the 600ms after sound onset and subtracting the mean deconvolved baseline activity in a 1 second interval before sequence onset^172^.

### Assessing single-neuron tuning to entropy

We quantified the degree to which single neurons were tuned to differences in entropy. To avoid stimulus-specific response features, we sorted our data according to sound identity, obtaining population response vectors (length equivalent to all sound-responsive neurons) for all presentations (i.e. trials) of each respective sound. These trials were pooled from different sequences and versions and therefore varied in entropy. For each sound and neuron, we computed the Spearman correlation between trial-wise sound-evoked activity and trial-wise entropy. This correlation provided a measure for how strongly a neuron’s activity tracked the information content of a given sound, or in other words, how strongly a neuron was tuned to entropy. We used Spearman rather than Pearson correlations to reduce sensitivity to outlying responses on individual trials. Repeating this procedure for all ten sounds, we obtained independent vectors expressing the correlations of neuronal activity to entropy for different sounds. To test how consistently neurons were tuned to entropy across sounds, we then correlated these vectors across all sounds.

### Representational similarity analysis to assess the population code of information content

We concatenated the sound-evoked population response vectors across all FOVs, and grouped the sound-evoked responses according to the entropy of single trials into five evenly sized bins (edges used for binning: [0.4395, 1, 1.5, 1.9183, 2.25, 3.4594] bits). We then built the mean population response vector for each sound and entropy bin, leading to 50 different response vectors (5 bins, 10 sounds). To assess how sound representations changed with entropy, we applied a representational similarity analysis^92,93,95,173^ by computing pairwise Pearson correlations among the 50 population response vectors. This yielded a 50×50 representational similarity matrix, with diagonal values of 1 by construction and off-diagonal elements reflecting the similarity between activity patterns evoked by different sound–entropy conditions. Based on this full representational similarity matrix, we then selected and displayed the 10×10 submatrices, expressing the representational similarities across all sounds in a given entropy bin.

### Display of representational maps

The representational relations of sound-evoked population responses under all entropy conditions are reflected by the 50×50 neuronal representational similarity matrix described above. However, because of the large number of pairwise correlations, this matrix is visually hard to interpret. Therefore, we generated a simplified display of the neuronal representational data as a low-dimensional *representational map*^92,95,174^. To obtain this representational map display, we performed a principal component analysis (PCA) on the full 50×50 representational similarity matrix. For the PCA, we used the built-in MATLAB function *pca* with a singular value decomposition. The resulting principal component scores of each sound and entropy condition served as a proxy of the representational pattern of each sound, relative to all other sounds. We plotted all ten sounds in each bin of entropy independently in the first two or three principal components, yielding a dimension-reduced display of the representational relations of all sounds in a given entropy. Comparing the PCA coordinates of given sounds across entropy bins revealed a systematic representational change with increasing information content.

### Characteristics of the representation of information content

To characterize the neural representation of information content in our experimental data, we quantified different features of the representational similarity matrices across entropy bins.

#### Increasing responsiveness over entropy

We quantified the extent to which increasing entropy was associated with stronger neuronal responses by computing the mean evoked ΔF/F_0_ response for each sound and entropy bin, and correlating this responsiveness magnitude vector (n=50 response magnitudes obtained from pooling responses of 10 sounds and 5 bins) with the entropy of each bin (using the mean entropy value of each element).

#### Consistency of representational stimulus relations over entropy

We quantified how stable the stimulus-to-stimulus relations were across entropy bins by taking the off-diagonal correlation pattern of the 10×10 representational similarity in each entropy bin, flattening it into a vector of 45 correlation values, and computing pairwise correlations between these vectors across entropy bins. This led to a 5×5 correlation matrix, expressing the similarity of representational patterns between sounds across the five entropy bins.

#### Mean representational similarity over entropy

We quantified the overall representational similarity in each bin of entropy. We pooled the 45 unique off-diagonal elements of each 10×10 similarity matrix across the five entropy bins (n=225 similarity values) and correlated these values with the corresponding mean entropy of each bin.

### Assessing representational axes of information content and overall responsiveness

We estimated the axis in representational space that best predicted entropy from the geometry of evoked population activity (see ref^91^ for an example application of this analytic approach). Specifically, we used a linear regression to fit the mean entropy of all 50 binned population response vectors (10 sounds, 5 entropy bins) with the 49 principal component scores from the representational map estimation (see above). The regression was conducted with a least-squares optimization and without intercept, yielding 49 component weights which together with the origin form the best linear fit of a particular sound feature in representational map space (for a display of this best-fitted entropy axis see Figure 4l or Supplementary Figure 7d). Since the lower-order principal components captured only a negligible amount of variance in representational similarities and could therefore potentially add noise to the axis estimate, we considered only the weights in the first 25 principal components for our subsequent analyses. We repeated this procedure for the mean response magnitude of all binned population response vectors, obtaining independent axes that reflect the encoded trajectory of entropy and response magnitude in representational space. We computed the congruence in the orientation of entropy and response magnitude axes by correlating their respective weights.

To estimate the range of correlations between the two axes expected from the observed data, we conducted a bootstrap analysis, fitting axes on re-sampled versions of the entropy and response magnitude data, respectively, obtained by randomly drawing data points with replacement. For each iteration of resampling the entropy and response magnitude data, we fitted the corresponding representational axes and computed their absolute Pearson correlation. Repeating this process 10,000 times, we obtained a distribution of absolute correlation values, reflecting the overall range of axis correlations in the data. We report the 95% confidence interval for this distribution and compare it to a value of 1 to demonstrate incomplete alignment between the axes for entropy and response magnitude.

### Regression of entropy from neuronal population activity and cross-stimulus generalization

Based on the representational axis of entropy, we quantified how strongly information content (i.e. entropy) was encoded in sound-evoked population activity, using a linear regression. We used the single-trial evoked population response vectors of a given sound (concatenated across all 5454 neurons), to train a regression model to predict the entropy of each respective trial. The regression was performed using the MATLAB function *fitrlinear* without intercept, providing a vector of regression weights for each neuron. We then tested cross-stimulus generalization by applying these fitted weights to predict the entropy of all single-trial population responses to a different sound and computed the resulting regression error (expressed as mean squared error). We repeated this procedure for all pairs of sounds used for training and testing, leading to a 10×10 error matrix that expressed how strongly a model trained on responses to one sound generalized to the decoding of information content from responses to other sounds. We compared the regression errors obtained from the data to analogous regressions where we shuffled the training data for each sound, randomly scrambling the entropy assignment across trials and the neuronal identities for each trial. For each training sound, we then quantified the mean squared regression error over all other predicted sounds. Based on the regression, for each neuron, we calculated the absolute mean regression weight across all sounds, providing a metric of how strongly single neurons contributed to the encoding of entropy on a population level which we refer to as the neuron’s “entropy weight”.

### Decoding entropy from neuronal population activity

To corroborate the regression of entropy based on neuronal population activity, we conducted a complementary decoding analysis. We used the single-trial evoked population response vectors of a given sound (concatenated across all 5454 neurons), and classified all trials into a “low entropy” or a “high entropy” class by separating at the median entropy. Based on this discretization of trials, we then trained a linear classifier (MATLAB function *lassoglm* with L1 regularization) to discriminate between the high and low entropy class. Cross-validation was performed using a leave-one-trial-out procedure; the classifier was trained on all but one trial and evaluated on the held-out trial. The mean decoding performance was then computed as the mean percentage of correctly classified trials for each sound.

### Decoding stimulus identity from neuronal population activity

To test the discriminability of sound-evoked neuronal activity patterns, we performed one-versus-all population decoding analyses using a linear classifier. Neural response data were arranged in a matrix with neurons as rows and stimulus presentations (trials) as columns. A corresponding vector indicated the identity of the sound presented on each trial. For each sound, trials of that sound were labeled as the positive class and trials of all other sounds were grouped as the negative class. Decoding was performed using a linear support vector machine (MATLAB function *fitclinear*, learner=SVM). Performance was evaluated using 5-fold cross-validation. In each fold, trials were randomly partitioned into training and test sets while preserving class labels. To avoid class imbalance during training, the training set was balanced by randomly subsampling the larger class so that equal numbers of target-sound trials and other-sound trials were used to fit the classifier. The trained classifier was then evaluated on the full held-out test set, which contained all remaining trials. Prior to classifier training, neural responses were z-scored across trials using the mean and standard deviation computed from the balanced training subset within each cross-validation fold, and the same transformation was applied to the corresponding test data. Decoding performance for each sound-versus-all comparison was quantified using balanced accuracy, defined as the mean of the true positive rate (fraction of correctly classified target-sound trials) and the true negative rate (fraction of correctly classified non-target trials).

To estimate the contribution of individual neurons to the discrimination of sounds, the absolute values of the linear classifier weights (syn. “identity weights”) were extracted from the trained model in each cross-validation fold and averaged across sounds.

### Comparing the encoding of entropy and stimulus identity

To test to what degree neurons contributing strongly to entropy encoding also contributed to stimulus-identity encoding, we correlated the log-transformed entropy weights and identity weights. To further assess how general response magnitude related to both measures, we conducted a linear regression of the absolute mean response of all neurons with their entropy and identity weights, using the MATLAB function *fitlm* with an intercept. Note that by considering the absolute value of the mean evoked response, we integrated excitatory and suppressive response patterns, providing a metric of overall change in neuronal activity relative to baseline.

### Comparison of the neural signatures of information content with simple repetition

To contrast the effects of entropy encoding with a simpler condition based on the mere number of previously sampled stimuli in a sequence, we replicated key analyses of representation for information content, binning responses according to the sounds’ positions in the sequence rather than entropy. We compared the resulting metrics reflecting the strength of the representational code to our analyses based on stimulus entropy (see Supplementary Figure 7f-h).

## Supporting information

Statistical Information

## Data availability

Original behavioral and neuronal data of this study have been deposited at G-Node (www.g-node.org) and will be made publicly available as of the date of publication. Any additional information related to the data is available from the corresponding authors upon reasonable request.

## Code availability

The original MATLAB code to analyze the data has been deposited at G-Node (www.g-node.org) and will be made publicly available as of the date of publication. Any additional information required to reanalyze the data reported in this paper is available from the corresponding authors upon reasonable request.

## Acknowledgements

We thank all participants who contributed to this study. We thank the team from the Mainz Behavioral and Experimental Laboratory (MABELLA), and Pascal Musial for the support during the data acquisition.

## Author contributions

S.R. and J.S. designed the study. J.S. conducted the human experiments. S.S. and J.S. analyzed the human EEG data. J.S. analyzed he human behavioral data. J.S. and L.W. conducted the behavioral experiments in mice. J.S. conducted the imaging experiments in mice. J.S. analyzed the mouse data. S.R. supervised the mouse experiments. O.T., F.M.D. and T.O.B. jointly supervised the EEG experiments. J.B.E. devised and performed the network model analyses. J.S. and S.R. wrote the manuscript. All authors edited the manuscript.

