## Supplementary Information for "Boredom and the representation of information content in the neocortex"

Supplementary Information (version 1)

### Supplementary Figures

#### Supplementary Figure 1

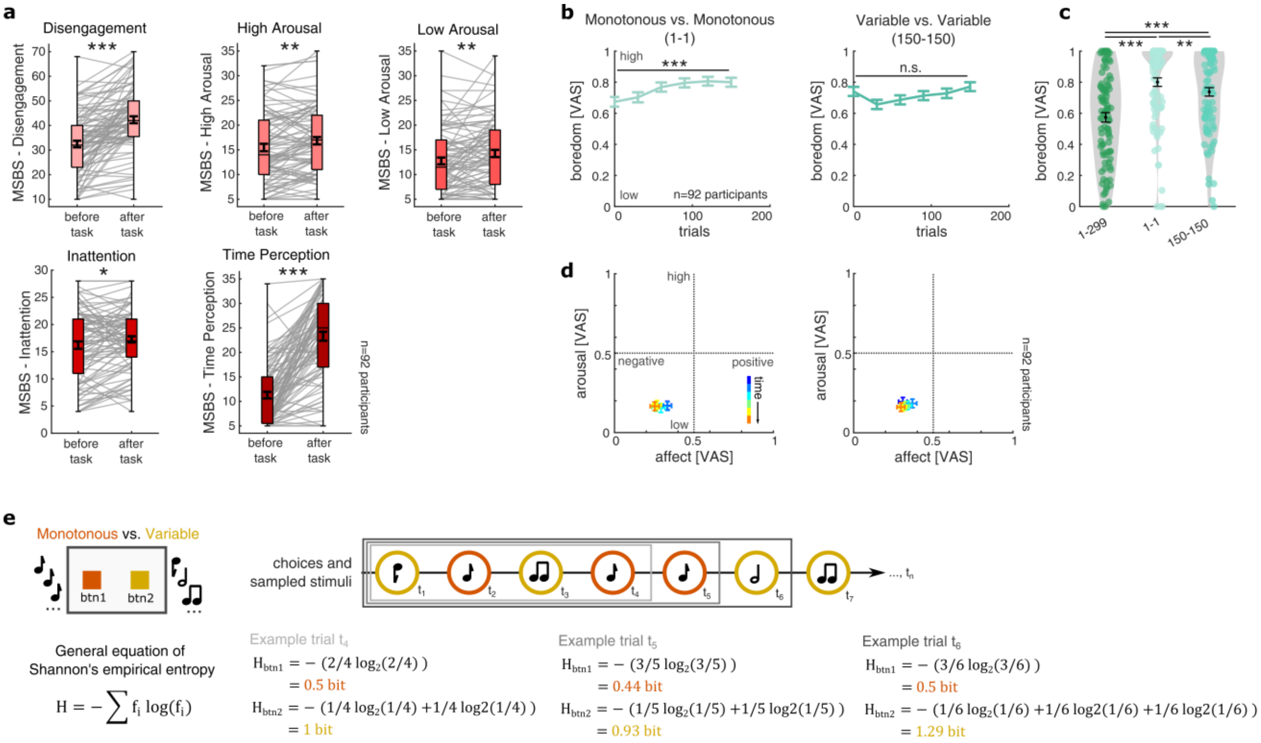


**Supplementary Figure 1 – Additional analyses of human choice task and sentiment:** (**a**) Multidimensional State Boredom Scale (MSBS) subscale scores before and after the choice task. All subscale scores increased after the task. Gray lines indicate individual participants; box plots show the median and interquartile range; thick black bars, mean ± SEM.; *: $p<0.05$, **: $p<0.01$, ***: $p<0.001$. (**b**) Boredom ratings increased over time in the mon-mon condition (left; **: $p<0.001$) but not in the var-var condition (right; n.s.: $p>0.05$). Data are mean ± SEM. (**c**) Mean state boredom across task conditions (n=92 participants), with highest ratings in the mon-mon (1-1) condition and lowest ratings in the mon-var (1-299) condition. Bars, mean ±SEM; **: $p<0.01$, ***: $p<0.001$. (**d**) Mean affect and arousal ratings over time in the mon-mon and var-var conditions. Data are mean ± SEM. (**e**) Schematic of trial-wise empirical entropy as a measure of experienced information content (Methods). We iteratively computed entropy at each trial for both alternatives, considering the probabilities of all stimuli sampled from a given alternative in previous trials relative to all stimuli sampled from both alternatives in previous trials. Repetitive sampling of the same stimulus in general led to decreasing entropy, whereas sampling of novel stimuli led to increasing entropy. This computation of entropy provides a metric expressing the diversity of previously sampled stimuli from each alternative which we used as a metric of experienced information content (see Methods for more details).

#### Supplementary Figure 2

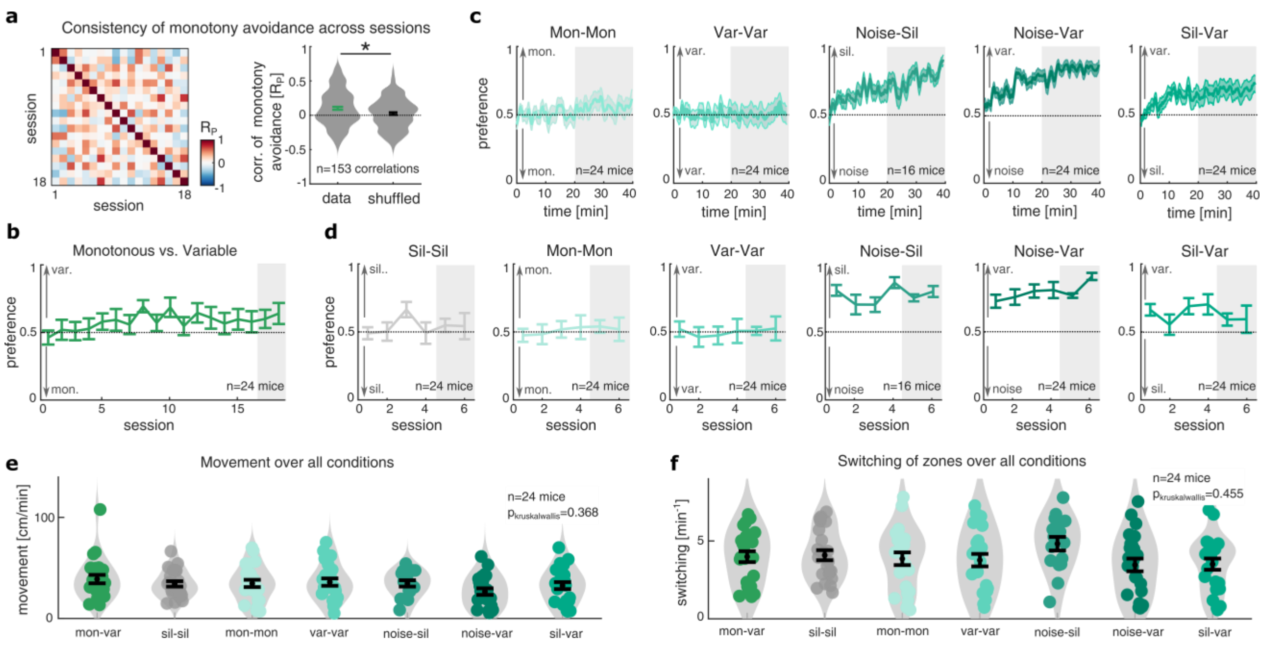


**Supplementary Figure 2 – Additional analyses of mouse place preference task:** (**a**) Since boredom has been described as both, a temporally limited and transient state, and a temporally stable trait expressing the general tendency to be bored, we wondered if monotony avoidance of individual mice could also reflect an individual trait. We hypothesized that if monotony avoidance would constitute a stable trait in individual mice, we would observe an overall positive correlation of monotony avoidance across sessions, whereas we expected no correlation across time in case of no trait components in the observed behavior. To test the stability of monotony avoidance in single mice across time, we computed the mean place preference for each mouse in the second half of the task for all 18 mon-var sessions that mice completed (left). This procedure provided us with a vector, expressing the monotony avoidance of all 24 mice on each session. We then computed the Pearson correlation matrix of monotony avoidance across sessions (n=24 mice). Comparing the correlations of monotony avoidance in our mice over time with a permuted version of the data, randomly shuffled across mice, we observed significant positive correlation of preference over sessions (right). *: $p<0.05$. This indicates a significant trait component in individual boredom-related behavior in mice. (**b**) Mean preference across mon-var sessions, showing increasing monotony avoidance over the first approximately eight sessions. Gray shading indicates the final two sessions detailed in Figure 2. Data are mean ± SEM. (**c**) Mean place preference over time in the control conditions. Gray shading indicates the second half used for quantification in Figure 2g. As expected, we find that mice show chance preference in conditions of equal stimulation at both zones, whereas they show preferential biases, avoiding loud noise and preferring variable sounds over complete silence, that build up over approximately 20 minutes in the task. Data are mean ± SEM. (**d**) Mean preference across sessions for the control conditions (as in b). Data are mean ± SEM. (**e**) Mean movement during the second half of each control condition, showing a large similarity of overall movement. Vertical bars are mean ± SEM. (**f**) Mean zone-switching frequency during the second half of each control condition (as in e). Mice show largely similar levels of overall switching across conditions. Vertical bars are mean ± SEM.

#### Supplementary Figure 3

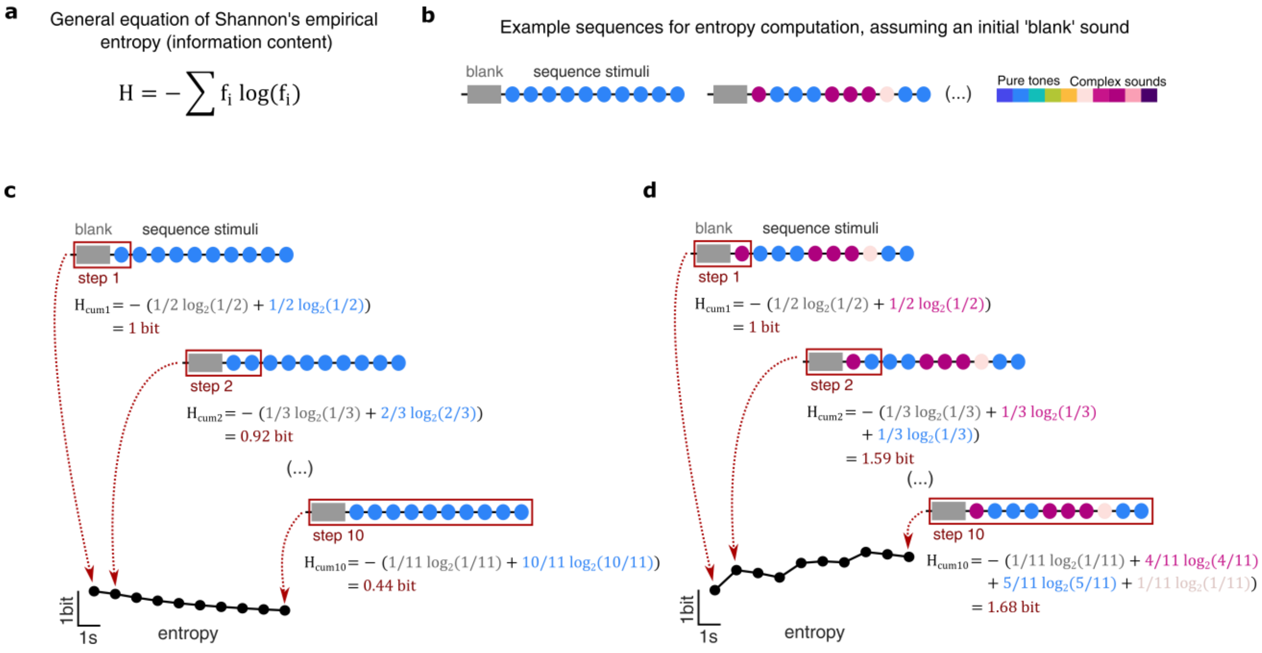


**Supplementary Figure 3 – Computing entropy as a measure of information content in sound sequences:** (**a**) General equation to compute entropy as a measure of information content, based on the relative occurrence probability of particular events, such as the occurrence of a sound stimulus (see Methods). (**b**) Two example sound sequences used in the experimentEach sequence comprised ten sounds drawn from a library of ten stimuli and was preceded by silence (see left sequence for a low diversity example, or right sequence for a higher entropy example; dots represent single sounds, color represents sound identity; here we use the color code corresponding to our mouse stimulus set, however, the same principles apply to the sequences used the human experiments (see Methods)). (**c,d**) Computation of entropy for example sequence with low or high entropy. To compute entropy, we considered the initial silence as an initial "blank" stimulus, and then computed the entropy for each of the following sounds, considering the relative probabilities of all stimuli in the sequence up to the current sound presentation. This procedure provides a metric of entropy that generally decreased when a stimulus is presented repetitively (see c), and increased when different stimuli are presented (see d). Entropy generally tended to increase with the number of sounds presented in a sequence. Maximum entropy was approached when many sounds occurred with similar frequencies.

#### Supplementary Figure 4

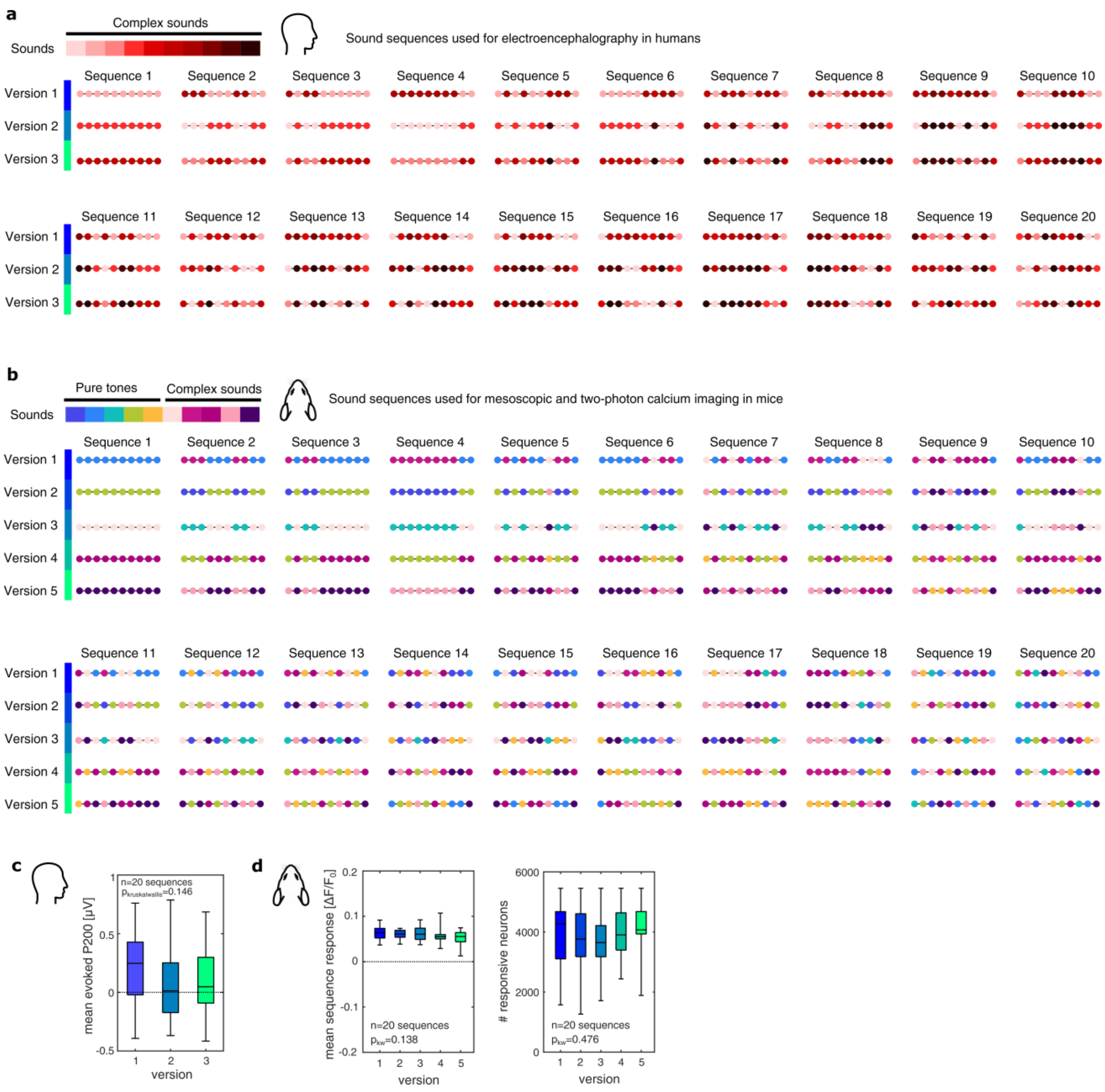


**Supplementary Figure 4 – Sound sequences with varying information content used in human and mouse experiments:** (**a**) Twenty sequence patterns used in human experiments, constructed from ten distinct 500-ms complex sounds. Sound identities are indicated by color. Three versions of each pattern were generated by reassigning sound identities while preserving sequence structure, reducing stimulus-specific biases. (**b**) The same 20 sequence patterns implemented for mouse experiments using ten distinct 70-ms sounds adapted to the mouse hearing range and previously shown to evoke distinct auditory cortical activity patterns (Aschauer et al., 2022, Noda et al., 2025). Five versions of each sequence pattern were generated. (**c**) Comparing the mean P200 component of the evoked EEG potentials to all sound sequences in each version, showed an overall comparability of responses across versions of the sequence. Box plots show the median and interquartile range. (**d**) Mean two-photon calcium response magnitude (left) and number of sound-responsive neurons (right) across mouse sequence versions showed widely comparable responses. Box plots show the median and interquartile range.

#### Supplementary Figure 5

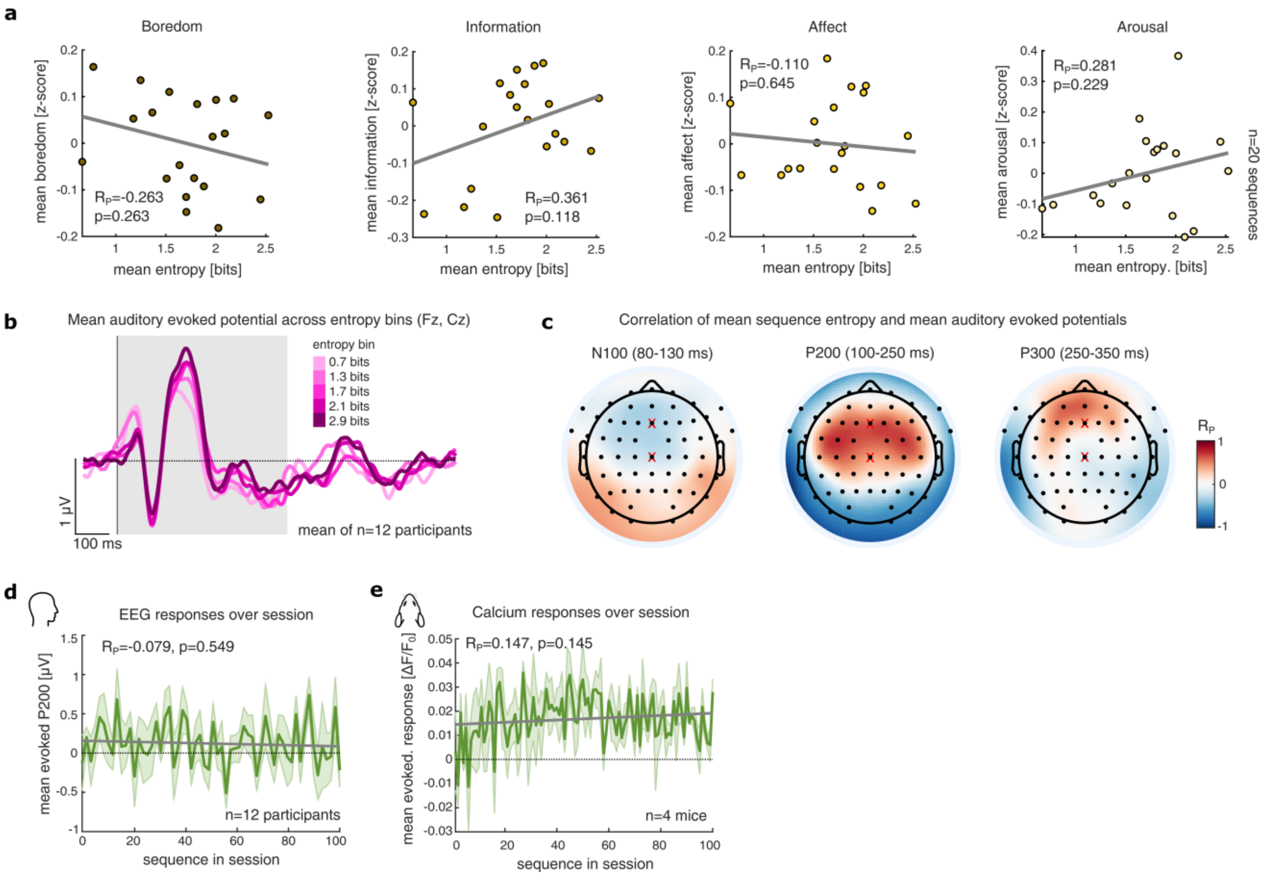


**Supplementary Figure 5 – Detailed sentiment of sound sequence and stable neural responses across EEG and mesoscopic imaging recordings:** (**a**) Correlations of mean sequence entropy and the mean sentiment ratings (average of 92 participants) across the 20 different sequence patterns. Gray lines show linear fits. The direction of the correlations matches the expected associations of information content and sentiment: Perceived boredom tended to decrease with growing entropy, perceived information content tended to increase with information content, and arousal tended to increase with growing entropy. (**b**) Mean sound-evoked EEG response across Fz and Cz, grouped into five entropy bins. The EEG response shows a strong modulation with entropy at a latency of approximately 200ms after stimulus onset. Shading indicates stimulus presentation; dashed line, baseline. (**c**) Scalp distributions of Pearson correlations between mean sequence entropy and mean N100, P200 and P300 amplitudes across 20 sequence patterns (equivalent analysis as in Figure 3e per electrode). The strongest association with entropy occurred for the P200 component at frontocentral and frontotemporal electrodes, consistent with expected auditory response patterns. Subsequent analyses therefore used mean P200 responses across Fz and Cz (red crosses). (**d**) Mean sound-evoked P200 responses remained largely stable throughout the imaging session. Note that the order of presenting the sound sequences was randomized. Data represented as mean ± SEM. (**e**) The mean sound-evoked mesoscopic calcium response also remained largely stationary throughout the experiment. Data represented as mean ± SEM.

#### Supplementary Figure 6

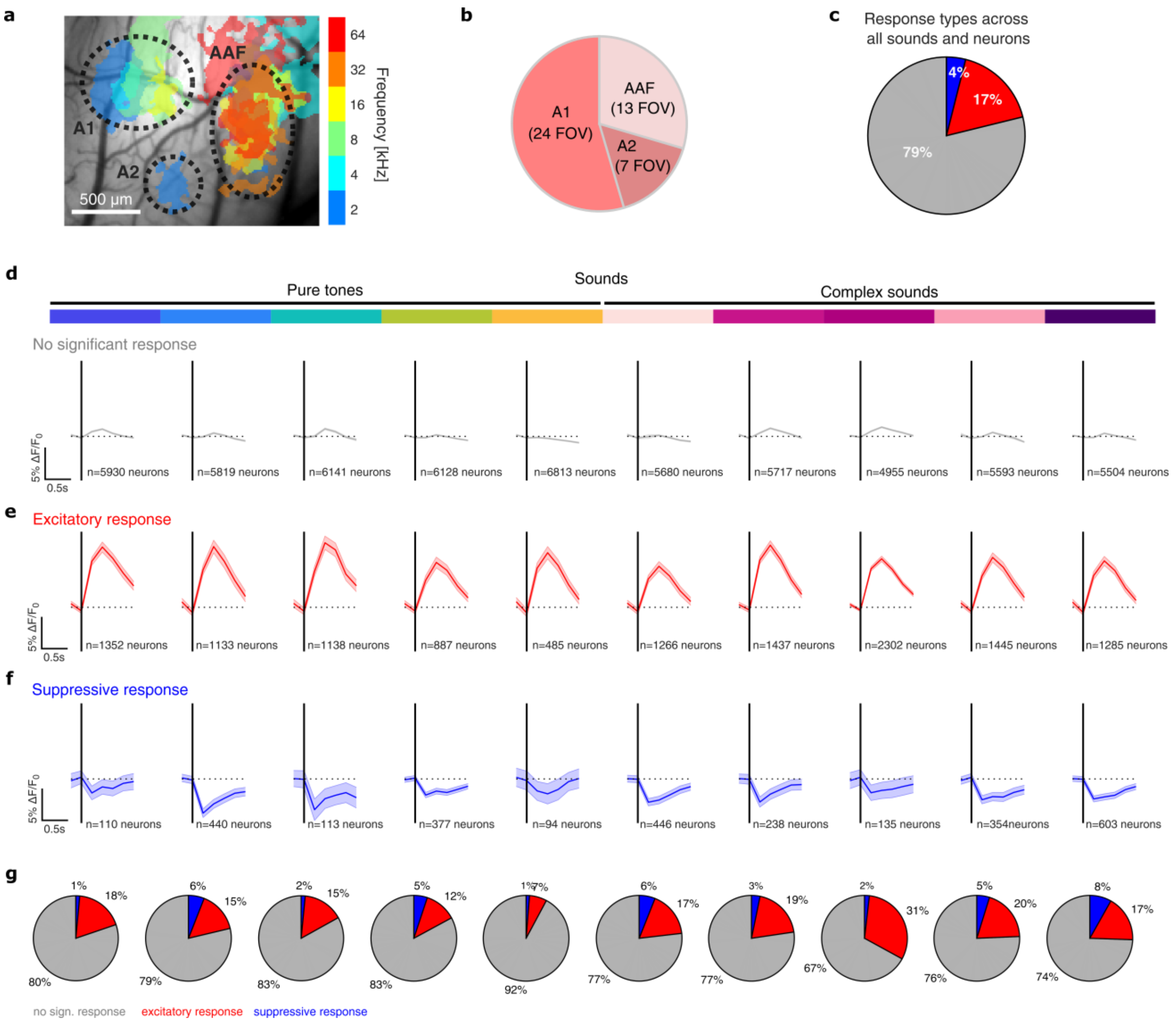


**Supplementary Figure 6 – Intrinsic imaging of the auditory cortex and neuronal response profiles to different sounds:** (**a**) Example cranial window and intrinsic imaging responses to pure-tone trains, showing the expected tonotopic organization of primary auditory cortex (A1), secondary auditory field (A2) and anterior auditory field (AAF). (**b**) Distribution of auditory fields across all FOV imaged in the two-photon experiment. (**c**) Distribution of sound-evoked calcium responses across all neurons and sounds (*n* = 73,920 responses from 7,392 neurons and 10 sounds). Responses were classified as non-significant (gray), excitatory (red) or suppressive (blue). (**d-f**) Mean evoked $\Delta F/F_0$ traces for non-significant (d), excitatory (e) and suppressive (f) responses across all individual sounds. Data are mean ± SEM. (**g**) Distribution of response classes for all sounds separately, illustrating qualitatively comparable response patterns across sounds. Some sounds deviated from the average fractions of response classes (e.g. see pure tone 5 and complex sound 3), so that we based our following analyses on within-sound comparisons of response patterns.

#### Supplementary Figure 7

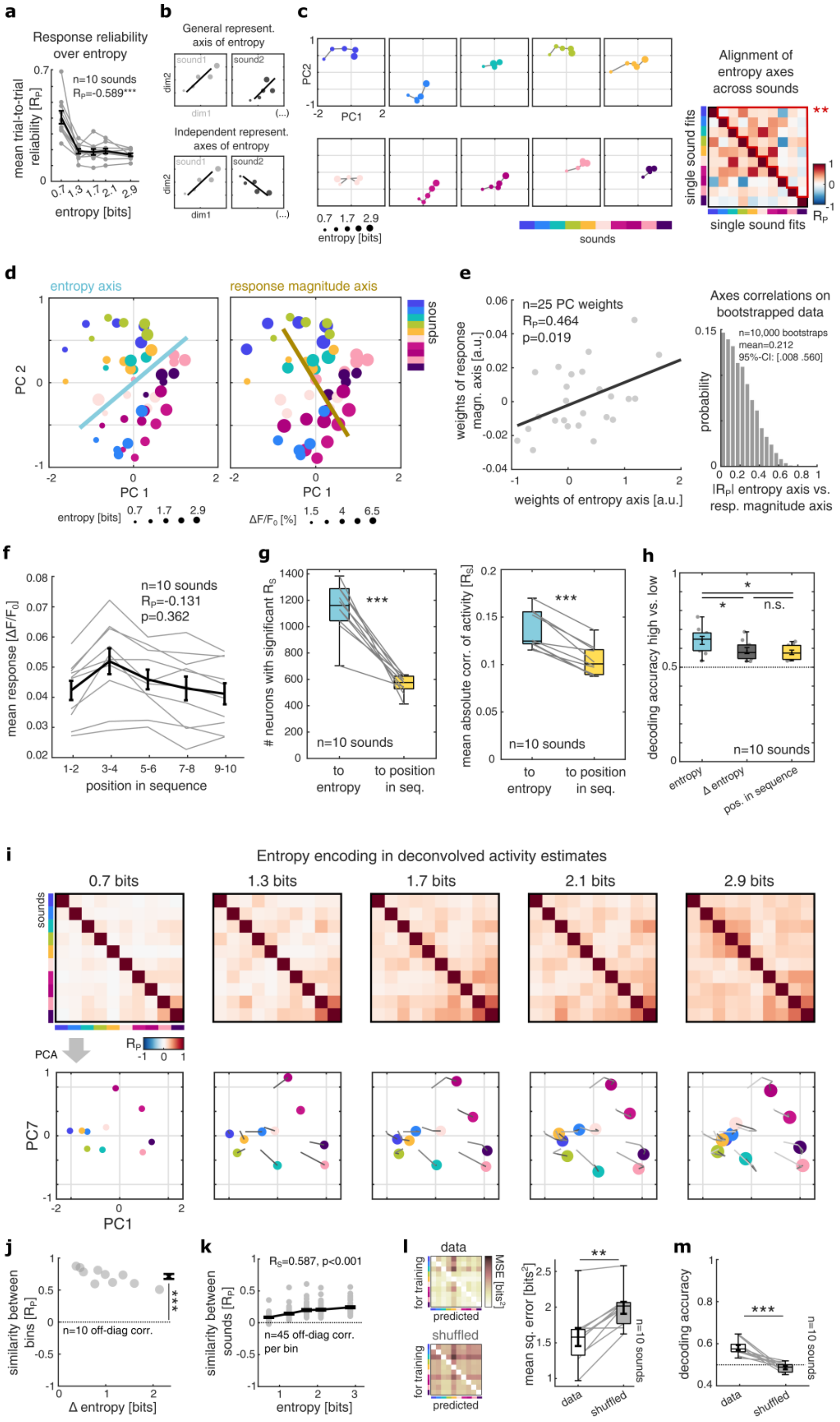


**Supplementary Figure 7 – The neural representation of information content is stimulus-invariant and robust against comparisons with representations of simple measures of stimulus repetition and deconvolution:** (**a**) Mean reliability of sound-evoked response patterns over entropy, computed as the average Pearson correlation between population response vectors across all sound-responsive neurons in each entropy bin. The initial drop indicates a higher reliability of responses to stimuli with lower entropy, typically associated with previous presentations of the same stimulus. (**b**) Scheme illustrating how representational map projections would be expected in case of (1) a general, stimulus-invariant representational code of information content where the representational axis of entropy would be shared across stimuli, or (2) a stimulus-specific representation of entropy, where each stimulus on a representational map has an independent axis of entropy. (**c**) PCA projections of sound representations across entropy bins (left). Colors indicate sound identity, point size entropy and gray lines neighboring entropy bins. Similar orientations across sounds indicate a shared entropy axis. Pairwise correlations between entropy axes fitted separately to each sound in the first 25 principal components are shown at right. Mean off-diagonal correlations were positive (*: $p<0.05$), supporting hypothesis 1 of a stimulus-invariant representation of information content. (**d**) PCA projection of sound representations, with point size indicating entropy (left) or mean response magnitude (right). Lines show fitted axes for entropy (blue) and response magnitude (ochre). Their incomplete alignment indicates that the population representation of entropy was not fully explained by response magnitude. Note that the left panel is equivalent to an overlayed display of Figure 4i, and that the displayed points and axes also correspond to a two-dimensional display of Figure 4l). (**e**) Correlation between weights of the entropy and response-magnitude axes across the first 25 principal components (left). To test what maximal and minimal degree of correlation could be expected between the entropy and response magnitude axes, we conducted a bootstrap analysis, sampling randomly with replacement from the distributions of entropy and response magnitude, respectively, fitting the axes on the resampled data, and computing their absolute Pearson correlation (right). We repeated this process 10,000 times, obtaining a distribution of axes correlations for the bootstrapped data, which we compared to a correlation value of 1. We observed significantly lower absolute correlations between the axes (mean $R_P$=0.22, 95%-CI = [0.008 0.560]), that despite an overall similarity, the representational axes of entropy and response magnitudes were distinct. (**f**) To compare in how far a neuronal representation of information content could also be explained by simpler stimulus features other than empirical entropy, we analyzed the neural response patterns as a function of position in the sequence. This approach assumed a general adaptation of evoked responses over the stimuli presented in a sequence. Analyzing the mean overall sound-evoked activity over stimulus positions did not show a significant correlation (n=10 sounds in 5 position bins), differing from entropy (see Figure 4d). Gray lines indicate individual sounds; box plots show the median and interquartile range. (**g**) Number of neurons significantly tuned to entropy or sequence position for each sound (left) and mean Spearman correlation between activity and either variable across sound-responsive neurons (n=5,454; right). Gray lines indicate individual sounds; box plots show the median and interquartile range. Panels f and g together indicate a stronger neuronal tuning to entropy than to simple stimulus repetitiveness. (**h**) Cross-validated decoding of entropy, trial-to-trial entropy change (Δentropy) and sequence position from population activity. Trials for each sound were classified into high and low categories for each variable. Overall, entropy was more strongly encoded than the change in entropy or position in the sequence. (**i-m**) Replication of population-level analyses using deconvolved neural activity (see Methods, compare the data plotted here with Figure 4 i-m). In general, we replicated the finding of a stimulus-invariant representation of information content, albeit slightly less pronounced (see i). The representational similarity matrices obtained from deconvolved data showed consistent structure across entropy bins (see j), were associated with increasing similarity over entropy (see k), and allowed a significant regression (see l) and decoding of stimulus entropy (see m). Together, these control analyses indicate that the neural representation of entropy, observed in our data, is robust against comparison to simpler metrics of stimulus repetition, is not fully explained by changes in evoked neural response magnitudes, and is robust against deconvolution.

#### Supplementary Figure 8

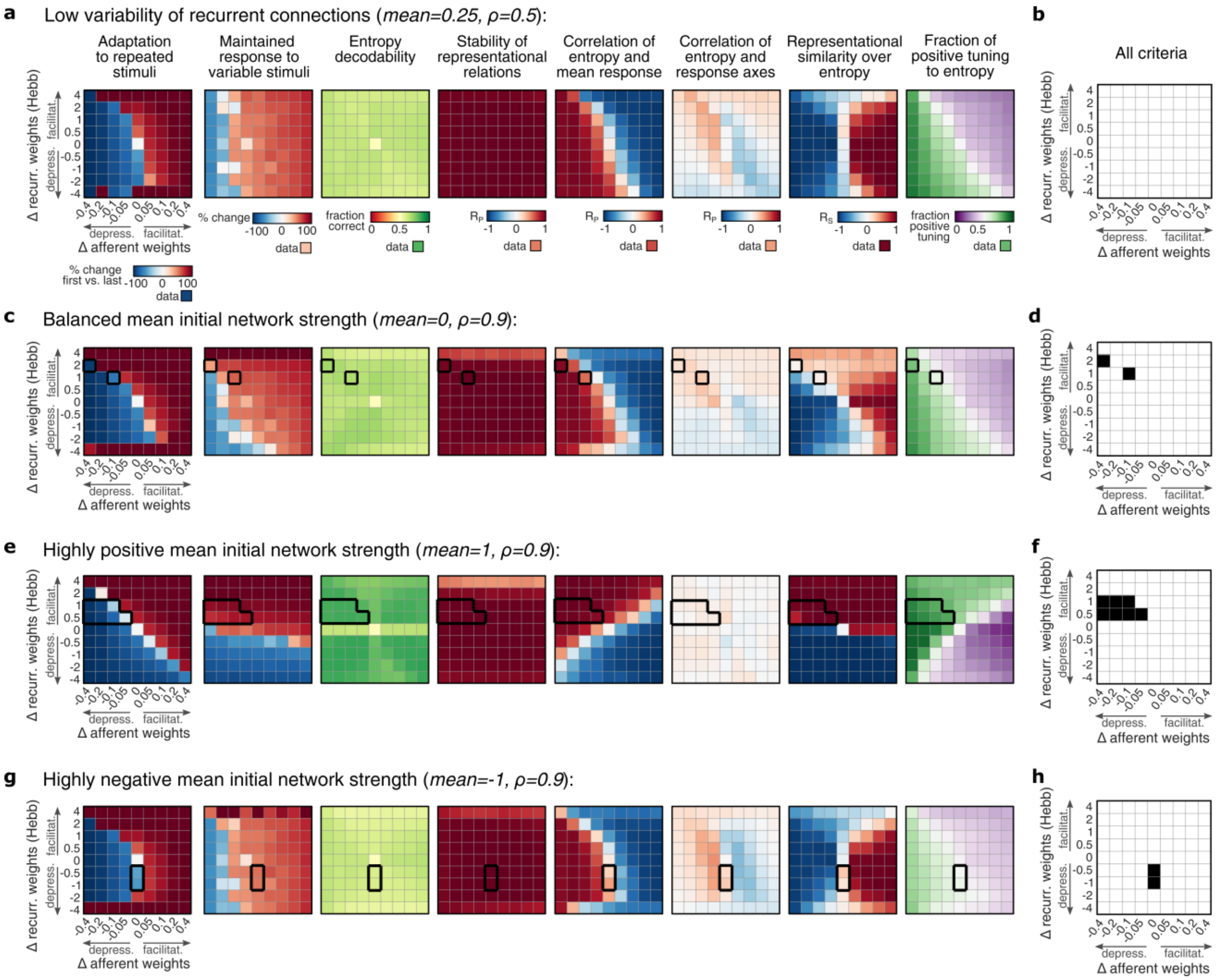


**Supplementary Figure 8 – Modeling a representation of information content in networks with varying initial network strength:** Based on the parameter scans for the two adaptation mechanisms of our network model, the change in recurrent weights (Δ recurrent) and the change in afferent gain (Δ afferent) (see Figure 5), we wondered how the ability to form a representation of information content was moderated by initial network strength (i.e. initial recurrent connectivity). To this end, we used the initial network parameters used for the simulations shown in Figure 5 (mean=0.25, ρ=0.9, reflecting a moderately excitatory network; see Methods) as a starting point, and systematically varied the initial weight distributions before repeating the parameter space analysis (number of simulations and other parameters are kept fixed, see Methods). (**a**,**b**) We first lowered the standard deviation of recurrent connections (ρ=0.5) to reduce the overall effects of recurrent connectivity in the model. Evaluation using the eight criteria (shown in a), adaptation to repeated stimuli, maintained responses for varying stimuli, entropy decodability, stability of representational relations, correlation of entropy and mean response, correlation of the entropy and response axes, representational similarity over entropy, and fraction of positive tuning to entropy, showed no working regime across the full parameter space (see fully blank matrix in b, working regime would be shown in black). This indicates that diverse recurrent connections in the network are essential to allow the formation of a representation of information content. (**c**,**d**) Same analyses for networks with balanced initial recurrent connections (mean=0, reflecting even fractions of excitatory and inhibitory connections in the network). Here, a working regime emerged under afferent depression and recurrent facilitation, aligning with our base model endowed with modest excitatory connections, however to a markedly lower degree. (**e**,**f**). When increasing the network strength to a strongly excitatory network (mean=1), we observed a representation of information content emerge for a broadened set of parameter combinations. Under this network configuration, modest recurrent facilitation tolerated higher degrees of afferent depression, still producing a working regime. (**g,h**) In an overall inhibitory network (mean=-1), we also observed a small working regime for mild recurrent depression without any afferent adaptation, which however was less pronounced than the working regimes in excitatory networks. Together, these analyses corroborate that afferent synaptic depression combined with recurrent synaptic facilitation in an excitatory network are most efficient in allowing the formation of a representation of information content of incoming stimuli.

### Supplementary Tables

#### Supplementary Table 1

| Conditions in choice task | 1-1 | 150-150 | 1-299 | Psychometric rating of sound sequences |
|---|---|---|---|---|
| # participants | 92 | | | |
| Participant IDs | 301-392 | | | |
| # sessions per condition | 1 | | | |
| Age (years) mean ± SD | 23.52 ± 3.17 | | | |
| Men / Women | 27 (29.4%) / 65 (70.6%) | | | |
| Right-handed / left-handed | 92 (100%) / 0 (0%) | | | |

**Supplementary Table 1 – Structure of human behavioral dataset**.

186 **Supplementary Table 2**

| Condition in place preference task | sil-sil | sil-noise | noise-83 | 1-1 | 41-41 | sil-83 | 1-82 |
|---|---|---|---|---|---|---|---|
| # mice | 24 | 16 | 24 | 24 | 24 | 24 | 24 |
| Mouse IDs | 205-212, 2083-2098 | 2083-2098 | 205-212, 2083-2098 | 205-212, 2083-2098 | 205-212, 2083-2098 | 205-212, 2083-2098 | 205-212, 2083-2098 |
| # sessions per condition | 6 | 6 | 6 | 6 | 6 | 6 | 20 |

187 **Supplementary Table 2 – Structure of mouse behavioral dataset**.

#### Supplementary Table 3

| Experiment | EEG recording | Pupillometry (a subset of EEG dataset) |
|---|---|---|
| # participants | 12 | 8 |
| Participant IDs | 401,402,403,404,405,406,407,451,452,454,455,456 | 401,402,403,404,405,406,407,454 |
| Age (years) mean ± SD | 24.92 ± 3.58 | 23.00 ± 1.93 |
| Men / Women | 4 (33.3%) / 8 (66.7%) | 1 (12.5%) / 7 (87.5%) |
| Right-handed / left-handed | 10 (100%) / 0 (0%) | 8 (100%) / 0 (0%) |

**Supplementary Table 3 – Structure of human EEG and pupillometry dataset**

#### Supplementary Table 4

| Data set (regressors) | n individuals with full data | n pooled trials across individuals, sequences and sets | Explained variance $R^2$ | F-statistic vs. constant model / p-value | Standard. regression weights [a.u.] | T-statistic of regressors | p-value of regressors |
|---|---|---|---|---|---|---|---|
| Mouse: calcium, pupil, movement | 3 | 3,000 | 0.0511 | 46.6 / $1.5*10^{-29}$ | 0.109<br>0.211<br>0.045 | 5.435<br>9.721<br>2.121 | $6.0*10^{-8}$<br>$5.8*10^{-22}$<br>0.034 |
| Human: EEG P200, pupil | 8 | 4,800 | 0.0024 | 5.66 / 0.003 | 0.047<br>0.015 | 3.232<br>1.003 | 0.001<br>0.316 |

**Supplementary Table 4 – Linear regression of entropy in single trials from mouse and human data.** The order of the reported regression weights corresponds to the order of regressors in the first column.

#### Supplementary Table 5

| Experiment | Mesoscopic imaging + pupillometry + movement | | | | |
|---|---|---|---|---|---|
| # mice | 5 | | | | |
| Mouse IDs | 1954 | 1955 | 1960 | 1962 | 1963 |
| Mesoscopic data | - | + | + | + | + |
| Pupillometry data | + | + | + | + | - |
| Movement data | + | + | + | + | + |

**Supplementary Table 5 – Structure of mouse mesoscopic imaging dataset**

#### Supplementary Table 6

| Experiment | Two-photon imaging | | | | | |
|---|---|---|---|---|---|---|
| Number of mice | 5 | | | | | |
| Mouse IDs | 1954 | 1955 | 1960 | 1962 | 1963 | **total** |
| # FOV per mouse | 11 | 10 | 7 | 6 | 10 | 44 |
| # good quality cells per FOV (mean ± SD) | 169 ± 22 | 170 ± 23 | 183 ± 21 | 176 ± 29 | 150 ± 20 | 168 ± 24 |
| # sound-responsive cells per FOV (mean ± SD) | 128± 44 | 135 ± 42 | 115 ± 40 | 125 ± 46 | 114 ± 25 | 123 ± 38 |
| # good quality cells total | 1854 | 1702 | 1279 | 1056 | 1501 | 7392 |
| # good quality sound-responsive cells total | 1403 | 1345 | 808 | 763 | 1135 | 5454 |

**Supplementary Table 6 – Structure of mouse two-photon imaging dataset**

#### Supplementary Table 7

| REAGENT OR RESOURCE | SOURCE | IDENTIFIER |
|---|---|---|
| **Bacterial and virus strains** | | |
| rAAV (ITR-hSyn-GCaMP6m-WPRE-ITR) | Aschauer et al., 2022, Cell Reports | N/A |
| rAAV (ITR-hSyn-H2B::mCherry-WPRE-ITR) | Aschauer et al., 2022, Cell Reports | N/A |
| **Deposited data** | | |
| Human behavioral and psychometric data; Human EEG data; Mouse behavioral data; Mouse mesoscopic calcium imaging data; Mouse two-photon calcium imaging data | This study | Repository at www.g-node.org. Will be made publicly available as of the date of publication. |
| **Experimental models: Organisms/strains** | | |
| *Homo sapiens*: Healthy student participants | Recruited via the ORSEE online system (Greiner et al., 2015, Journal of the Economic Science Association) | N/A |
| *Mus musculus*: CB57BL/6J | Jackson Laboratory | Strain #000664; RRID: IMSR_JAX:000664 |
| **Recombinant DNA** | | |
| pAAV-hSyn-GCaMP6m-WPRE | Aschauer et al., 2022, Cell Reports | N/A |
| pAAV-hSyn-H2B::mCherry-WPRE | Aschauer et al., 2022, Cell Reports | N/A |
| **Software and algorithms** | | |
| MATLAB R2022a | MathWorks, Natick, MA, USA | N/A |
| Image processing of chronic two-photon data | Aschauer et al., 2022, Cell Reports | https://doi.org/10.5281/zenodo.5822486 |
| Processing and analysis of behavioral and neuronal data | This study | Repository at www.g-node.org. Will be made publicly available as of the date of publication. |

**Supplementary Table 7 – Key resource table**
