## Supplementary material for "Boredom and the representation of information content in the neocortex": Statistical Information

Statistical information (version 1)

| Main analyses | | | | | |
|---|---|---|---|---|---|
| **Figure** | **Compared metrics** | **Statistical Test** | **$n_1$** | **$n_2$** | **p-value, other metrics** |
| 1c | Mean preference in second half vs. 0.5 | t-test | 92 participants | - | mon-var: p<0.001<br>mon-mon: p=0.801<br>var-var: p=0.296 |
| 1d | MSBS sum score (boredom questionnaire) before and after the choice task | Wilcoxon signed rank test | 92 participants | 92 participants | p<0.001 |
| 1e | Correlation between mean boredom rating and mean preference in second half in mon-var | Pearson correlation | 92 participants | - | R=0.211, p=0.043 |
| 1f | Initial versus last boredom rating during the choice task | Wilcoxon signed rank test | 92 participants | 92 participants | p<0.001 |
| 1g | Initial versus last affect and arousal ratings during the choice task | Wilcoxon signed rank test | 92 participants | 92 participants | affect: p<0.001<br>arousal: p<0.001 |
| 1i | Mean entropy sampled from monotonous versus variable button | Wilcoxon signed rank test | 92 participants | 92 participants | p<0.001 |
| 1k | Fraction of correctly predicted choices by the logistic regression model vs. 0.5 | t-test | 92 participants | - | p<0.001 |
| 1l | Contributions of the model parameters: idiosyncratic bias, entropy difference, previous choice | Wilcoxon signed rank test | 92 participants | 92 participants | bias vs. entropy diff.: p<0.001<br>bias vs. prev. choice: p<0.001<br>entropy diff. vs. prev. choice: p=0.004 |
| 2g | Mean preference in second half vs. 0.5 | t-test | 24 mice | - | mon-var: p=0.015<br>sil-sil: p=0.239<br>mon-mon: p=0.263<br>var-var: p=0.866<br>noise-sil: p<0.001<br>noise-var: p<0.001<br>sil-var: p=0.004 |
| 2h | Mean entropy sampled from monotonous versus variable zone | Wilcoxon signed rank test | 24 mice | 24 mice | p<0.001 |
| 2j | Fraction of correctly predicted choices by the logistic regression model vs. 0.5 | t-test | 24 mice | - | p<0.001 |
| 2k | Contributions of the model parameters: idiosyncratic bias, entropy difference, previous choice | Wilcoxon signed rank test | 24 mice | 24 mice | bias vs. entropy diff.: p=0.007<br>bias vs. prev. choice: p=0.021<br>entropy diff. vs. prev. choice: p<0.001 |
| 3c | Mean ratings over the five sequences with the highest mean entropy versus the mean over the five | Paired one-sided t-test:<br>high entropy – low boredom; | 92 participants | 92 participants | boredom: p=0.018<br>information: p=0.045<br>affect: p=0.584<br>arousal: p=0.024 |

| | | | | | |
|---|---|---|---|---|---|
| | sequences with the lowest mean entropy | high entropy - high information;<br>high entropy - high/pos. affect;<br>high entropy - high arousal | | | |
| 3e | Correlation of mean sequence entropy and the mean EEG P200 potential or pupil response in 500ms after sound onset | Pearson correlation | 20 sequences | - | EEG: R=0.569, p=0.009<br>Pupil: R=0.542, p=0.014 |
| 3f | Regression of single trial entropy with mean EEG P200 potential and pupil response | Linear regression with intercept | 4,800 trials (pooled from 8 participants with 600 sound presentations per participant) | - | EEG: p=0.001<br>Pupil: p=0.316 |
| 3k | Correlation of mean sequence entropy and the mean mesoscopic calcium response, pupil response, or movement in 600ms after sound onset | Pearson correlation | 20 sequences | - | Calcium: R=0.465, p=0.039<br>Pupil: R=0.344, p=0.137<br>Movement: R=0.237, p=0.314 |
| 3l | Regression of single trial entropy with mean mesoscopic calcium response, pupil response and movement in 600ms after sound onset | Linear regression with intercept | 5000 trials (pooled from 5 mice with 1000 sound presentations per mouse) | - | Calcium: p<0.001<br>Pupil: p<0.001<br>Movement: p=0.034 |
| 4d | Correlation of the mean binned activity of all neurons and the entropy of each response bin | Pearson correlation | 50 data points (10 sounds over 5 entropy bins) | - | Pooled: R=0.363, p=0.010<br>Excitatory: R=0.326, p=0.021<br>Suppressive: R=0.012, p=0.937 |
| 4h | Correlations of entropy tuning of neurons across sounds versus 0 | t-test | 45 correlations (unique off-diagonal elements of correlation matrix with 10x10 sounds) | - | p<0.001 |
| 4j | Correlations of representational map pattern across entropy bins versus 0 | t-test | 10 correlations (unique off-diagonal elements of correlation matrix with 5x5 bins) | - | p<0.001 |

| | | | | | |
|---|---|---|---|---|---|
| 4k | Rank correlation of pairwise representational similarities and entropy | Spearman correlation | 225 data points (45 unique off-diagonal correlations for 5 entropy bins) | - | R=0.760, p<0.001 |
| 4m | Mean squared error of linear regression model of entropy from the real data versus shuffled data of each sound | Wilcoxon signed rank test | 10 sounds | 10 sounds | p=0.002 |
| 4n | Rank correlation between entropy weight identity weight | Spearman correlation | 5454 neurons | - | R=0.039, p=0.004 |
| 4o | Regression of the absolute mean calcium response of each neuron from its entropy weight and identity weight | Linear regression with intercept | 5454 neurons | - | Entropy weight: p<0.001<br>Sound identity weight: p<0.001 |
| 4o | Contributions of the parameters for absolute mean calcium responses: entropy regression weight versus sound identity decoding weight | Wilcoxon signed rank test | 5454 neurons | 5454 neurons | p<0.001 |

### Supplementary analyses

| Figure | Compared metrics | Statistical Test | $n_1$ | $n_2$ | p-value, other metrics |
|---|---|---|---|---|---|
| S1a | MSBS subscale scores before and after the choice task | Wilcoxon signed rank test | 92 participants | 92 participants | Disengagement: p<0.001<br>High Arousal: p=0.001<br>Low Arousal: p=0.009<br>Inattention: p=0.013<br>Time Perception: p<0.001 |
| S1b | Initial versus last boredom rating during the choice task | Wilcoxon signed rank test | 92 participants | 92 participants | Mon-Mon: p<0.001<br>Var-Var: p=0.057 |
| S1c | Mean boredom rating across the task conditions | Wilcoxon signed rank test | 92 participants | 92 participants | Mon-Var vs. Mon-Mon: p<0.001<br>Mon-Var vs. Var-Var: p<0.001<br>Mon-Mon vs. Var-Var: p=0.007 |
| S1d left | Initial versus last affect and arousal ratings during the Mon-Mon task | Wilcoxon signed rank test | 92 participants | 92 participants | affect: p<0.001<br>arousal: p=0.403 |
| S1d right | Initial versus last affect and arousal ratings during the Var-Var task | Wilcoxon signed rank test | 92 participants | 92 participants | affect: p=0.085<br>arousal: p=0.067 |
| S2a | Correlation of place preference of all mice in mon-var task across | Wilcoxon rank sum test | 153 (unique off-diagonal elements of | 153 (unique off-diagonal elements of | p=0.021 |

| | | | | | |
|---|---|---|---|---|---|
| | sessions, versus correlation shuffled across mice | | correlation matrix with 18x18 sessions, mean over 24 mice) | correlation matrix with 18x18 sessions, computed mean over 24 mice shuffled for each session) | |
| S2e | Mean movement in second half across task conditions | Kruskal-Wallis test | 24 mice | - | p=0.368 |
| S2f | Mean switching frequency in second half across task conditions | Kruskal-Wallis test | 24 mice | - | p=0.455 |
| S4c | Mean evoked auditory EEG P200 potential over all participants and sequences across different versions of the sequences | Kruskal-Wallis test | 20 sequences | - | p=0.146 |
| S4d left | Mean evoked two-photon calcium response over all mice and sequences across different versions of the sequences | Kruskal-Wallis test | 20 sequences | - | p=0.138 |
| S4d right | Mean number of sound-responsive neurons over all mice and sequences across different versions of the sequences | Kruskal-Wallis test | 20 sequences | - | p=0.476 |
| S5a | Correlation of mean sequence entropy and the mean psychometric ratings of boredom, information, affect and arousal of all sequences | Pearson correlation | 20 sequences | - | Boredom: R=-0.263, p=0.263<br>Information: R=0.361, p=0.118<br>Affect: R=-0.110, p=0.645<br>Arousal: R=0.281, p=0.229 |
| S5d | Correlation of mean sound evoked auditory P200 responses (over 12 participants) and the sequence counts in a session | Pearson correlation | 60 presented sequences in a session | - | R=-0.079, p=0.549 |
| S5e | Correlation of mean sound evoked mesoscopic calcium responses (over 4 mice) and the sequence counts in a session | Pearson correlation | 100 presented sequences in a session | - | R=0.147, p=0.145 |
| S7a | Correlation of entropy bin indices and the mean trial-to-trial reliability of response vectors for each sound and entropy bin | Pearson correlation | 50 data points (10 sounds over 5 entropy bins) | - | R=-0.589, p<0.001 |
| S7c | Pairwise correlations of the representational axes (in | t-test | 45 (unique off-diagonal elements of | - | p=0.001 |

| | | | | | |
|---|---|---|---|---|---|
| | PCA space) of entropy fitted for all sounds versus 0 | | correlation matrix with 10x10 sounds) | | |
| S7e left | Correlation of the representational axis (in PCA space) of entropy versus the axis of response magnitude | Pearson correlation | 25 weights for first principal components | - | R=0.464, p=0.019 |
| S7e right | Distribution of Pearson correlations between the axes, fitted on bootstrapped entropy and response magnitude data, and comparison to a correlation of 1 | Bootstrap test | Distribution of correlations for 10,000 bootstraps | - | P<0.001 |
| S7f | Correlation of the mean binned evoked activity of all neurons and the bin index (binned by position in sequence) | Pearson correlation | 50 data points (10 sounds over 5 position bins) | - | R=-0.131, p=0.362 |
| S7g left | Number of neurons with a significant rank correlation of evoked activity and entropy, or sound position in the sequence (after Bonferoni-Holm correction) | t-test | 10 sounds | 10 sounds | p<0.001 |
| S7g right | Mean rank correlation of evoked activity and entropy, and of evoked activity and sound position in the sequence | t-test | 10 sounds | 10 sounds | p<0.001 |
| S7h | Decoding accuracy for responses classified into high or low categories based on entropy, Δ entropy, or sound position in the sequence versus 0.5 (chance level) | t-test | 10 sounds | - | entropy: p<0.001<br>Δ entropy: p<0.001<br>position in seq.: p<0.001 |
| S7h | Decoding accuracy for responses classified into high or low categories based on entropy, Δ entropy, or sound position against each other | t-test | 10 sounds | 10 sounds | entropy vs. Δ entropy: p=0.037<br>entropy vs. pos. in seq.: p=0.049<br>Δ entropy vs. pos. in seq.: p=0.625 |
| S7j | Correlations of representational map pattern across entropy bins versus 0 (obtained from deconvolved data) | t-test | 10 correlations (unique off-diagonal elements of correlation matrix with 5x5 bins) | - | p<0.001 |
| S7k | Rank correlation of pairwise representational similarities of deconvolved data and entropy | Spearman correlation | 225 data points (45 unique off-diagonal | - | R=0.587, p<0.001 |

| | | | | | |
|---|---|---|---|---|---|
| | | | correlations for 5 entropy bins) | | |
| S7l | Mean squared error of linear regression model of entropy from the normal deconvolved data versus shuffled deconvolved data of each sound | Wilcoxon signed rank test | 10 sounds | 10 sounds | p=0.002 |
| S7m | Decoding accuracy of high vs. low entropy bins from the deconvolved data | Wilcoxon signed rank | 10 sounds | 10 sounds | p<0.001 |

**Statistical analyses -** All statistical tests were two-sided unless otherwise specified.
